# Conserved FimK truncation coincides with increased expression of type 3 fimbriae and cultured bladder epithelial cell association in *Klebsiella quasipneumoniae*

**DOI:** 10.1101/2022.04.27.489788

**Authors:** Sundharamani Venkitapathi, Yalini H. Wijesundara, Fabian C. Herbert, Jeremiah J. Gassensmith, Philippe E. Zimmern, Nicole J. De Nisco

## Abstract

*Klebsiella spp*. commonly cause both uncomplicated urinary tract infection (UTI) and recurrent UTI (rUTI). *Klebsiella quasipneumoniae*, a relatively newly defined species of Klebsiella, has been shown to be metabolically distinct from *Klebsiella pneumoniae*, but its urovirulence mechanisms have not been defined. *K. pneumoniae* uses both type 1 and type 3 fimbriae to attach to host epithelial cells. The type 1 fimbrial operon is well-conserved between *Escherichia coli* and *K. pneumoniae* with the exception of *fimK*, which is unique to *Klebsiella spp*. FimK contains an N-terminal DNA binding domain and a C-terminal phosphodiesterase (PDE) domain that has been hypothesized to cross-regulate type 3 fimbriae expression via modulation of cellular levels of cyclic di-GMP. Here, we find that a conserved premature stop codon in *K. quasipneumoniae fimK* results in truncation of the C-terminal PDE domain and that the bladder epithelial cell association and invasion of uropathogenic *K. quasipneumoniae* strain KqPF9 is dependent on type 3 but not type 1 fimbriae. Further, we show that basal expression of both type 1 and type 3 fimbrial operons as well as bladder epithelial cell association is elevated in KqPF9 relative to uropathogenic *K. pneumoniae* TOP52 and that complementation of KqPF9*ΔfimK* with the TOP52 *fimK* allele reduced type 3 fimbrial expression and bladder epithelial cell attachment. Taken together these data suggest that the C-terminal PDE of FimK can modulate type 3 fimbrial expression in *K. pneumoniae* and its absence in *K. quasipneumoniae* may lead to a loss of type 3 fimbrial cross-regulation.

**Importance:** *K. quasipneumoniae* is often indicated as the cause of many opportunistic infections including urinary tract infection (UTI), which affects >50% of women worldwide. However, virulence mechanisms of *K. quasipneumoniae* remain uninvestigated. Prior to this work, *K. quasipneumoniae* and *K. pneumoniae* had only been distinguished phenotypically by metabolic differences. This work contributes to the understanding of *K. quasipneumoniae* virulence phenotypes by evaluating the contribution of type 1 and type 3 fimbriae, which are critical colonization factors encoded by all *Klebsiella spp*., to *K. quasipneumoniae* bladder epithelial cell attachment. We identify clear phenotypic differences in both bladder epithelial cell attachment between uropathogenic *K. pneumoniae* and *K. quasipneumoniae*. Importantly, we find that a structural difference in the fimbrial regulatory gene *fimK* may contribute to differential co-regulation of type 1 and type 3 fimbriae between the two *Klebsiella* species.

## Introduction

*Klebsiella spp*. including *K. pneumoniae, K. variicola, and K. quasipneumoniae* are common causes of both acute and recurrent UTI (1-5). Recurrent UTI (rUTI), defined as two symptomatic UTI episodes within six months or three within twelve months, poses a major health issue with ∼50% of UTIs in postmenopausal women estimated to develop into rUTI (6-8). Uropathogenic *E. coli* (UPEC) is the most common organism implicated in rUTI, and is followed in prevalence by the genus *Klebsiella*, which accounts for 15-17% cases (9, 10). *K. pneumoniae* has also been reported to be one of the most common causes of hospital acquired UTI (11, 12). *K. pneumoniae* and closely related species *K. quasipneumoniae* and *K. variicola* are a growing clinical concern due to the prevalence of multi-drug resistant strains in patients with both community-acquired, and hospital-acquired UTI (2, 3). Alongside extended spectrum beta-lactamases (ESBLs), *Klebsiella spp*. isolated from UTI patients often encode carbapenemases, aminoglycoside-modifying enzymes, and fluoroquinolone resistance genes (13-17).

*K. pneumoniae* expresses both type 1 and type 3 fimbrial operons (18, 19). These fimbriae have been reported to play a role in attachment and invasion to bladder epithelial cells in a mannose sensitive and insensitive fashion, respectively (20-23). Type 1 fimbriae are encoded by the *fim* operon and are expressed among most *Enterobacteriaceae* (24, 25). The *fim* operon of *K. pneumoniae* shares a high degree of similarity with *E. coli* (26) except for the presence of *fimK* at the end of the operon (27). The expression of type 1 fimbriae is regulated by phase transition of the invertible *fimS* regulatory sequence (*fim* switch) (28). The phase variation is mediated by recombinases FimB and FimE that are also encoded within the *fim* operon (29). Type 3 fimbriae, though initially identified in *Klebsiella spp*., have also been reported in other *Enterobacteriaceae* including *Serratia spp*., *Enterobacter spp*. and more recently encoded within plasmids of a few *E. coli* isolates (30-32). Type 3 fimbriae are encoded by the *mrk* operon with *mrkA* encoding the major structural subunit and *mrkD* encoding the adhesin (33, 34). The expression of type 3 fimbriae is regulated by the *mrkHIJ* gene cluster (35, 36). MrkH is a PilZ domain containing protein that functions as the major activator of *mrkABCDF* transcription upon binding to the second messenger cyclic-di-GMP. MrkI has been reported to contain a LuxR type DNA binding domain and functions as a minor activator of *mrkABCDF* (36). MrkJ acts as a phosphodiesterase (PDE) hydrolyzing cyclic di-GMP preventing its binding to MrkH and thereby repressing *mrkABCDF* expression (35).

*K. quasipneumoniae* were previously classified as part of *K. pneumoniae* phylogroups KpIIA and KpIIB but were relatively recently distinguished from *K. pneumoniae* as a new species with distinct metabolic phenotypes (37). *K. quasipneumoniae* isolates have been collected from patients with bloodstream infections as well as from the urine of patients with uncomplicated UTI and rUTI (3, 5, 38). However, little is known about how *K. quasipneumoniae* interacts with the bladder environment. In this study, genomic analysis *K. quasipneumoniae* showed a difference in the structure of the type 1 fimbrial regulatory gene *fimK* compared to *K. pneumoniae*. The *K. quasipneumoniae fimK* allele is truncated and lacks the C-terminal PDE domain conserved among *K. pneumoniae* isolates. Because the PDE activity of *K. pneumoniae* FimK has been hypothesized to cross-regulate type 3 fimbriae by reducing cyclic-di-GMP levels necessary for MrkH activation, we sought to investigate differences in type 3 fimbrial regulation between the two species (39). We show that attachment of *K. quasipneumoniae* strain KqPF9 to cultured bladder epithelial cells is dependent on type 3 but not type 1 fimbriae. Further, we demonstrate that the *fimK* C-terminal PDE domain downregulates type 3 fimbriae when expressed in both *K. pneumoniae* and *K. quasipneumoniae*, but that *K. quasipneumoniae fimK*, which lacks this domain, does not affect type 3 fimbriae expression. Taken together, this study defines the contribution of type 1 and type 3 fimbriae to *K. quasipneumoniae* bladder epithelial cell association and identifies a possible role for *fimK* in regulation of type 3 fimbriae in uropathogenic *K. pneumoniae* but not in *K. quasipneumoniae*.

## Results

### Attachment and invasion of uropathogenic *Klebsiella quasipneumoniae* to human bladder epithelial cells is mannose insensitive

Cell association and invasion phenotypes of *Klebsiella quasipneumoniae* have not been previously reported. We therefore first sought to use KqPF9, an uropathogenic *K. quasipneumoniae* strain that we recently isolated from a postmenopausal woman with rUTI to evaluate *K. quasipneumoniae* cell association and invasion. The complete genome of KqPF9 has been previously reported and contains one chromosome (5.27 Mbp) and 4 plasmids (of sizes 399,394 bp; 4,730 bp; 4,096 bp and 4,000 bp) (5). Because analysis of the KqPF9 genome revealed that the chromosome encodes both type 1 (*fim*) and type 3 (*mrk*) fimbrial operons, we first sought to determine if KqPF9 bladder epithelial cell association was mannose dependent, which would suggest that it was predominantly mediated by type 1 fimbriae (5, 40) (Figure 1A, 1B). We measured the effect of D-mannose on KqPF9 association with bladder epithelial cell line 5637 (ATCC) via cell association assay using *K. pneumoniae* 78578 (KpMGH78578, ATCC), which expresses both type 1 and type 3 fimbriae, as a mannose insensitive control and uropathogenic *E. coli* UTI89, which expresses only type 1 fimbriae, as a mannose sensitive control (19, 40-42). While a significant 75.4% reduction in 5637 bladder epithelial cell association was observed for UTI89 in the presence of D-mannose, KqPF9 and KpMGH78578, showed no significant decrease in association (Figure 1C). Then, using gentamicin protection assays to measure bladder epithelial cell invasion, we observed that D-mannose treatment similarly reduced invasion frequencies of UTI89 (56.3%), while KqPF9 and KpMGH78578 invasion frequencies were not significantly reduced (Figure 1D). This pattern of invasion is consistent with previous reports in uropathogenic *K. pneumoniae* isolate 3091 that expresses both type 1 and type 3 fimbriae (23). These results suggest that as in *K. pneumoniae*, adhesion and invasion of bladder epithelial cells by *K. quasipneumoniae* KqPF9 is mannose-independent and may not rely on type 1 fimbriae. To confirm these results, we also evaluated mannose sensitivity of KqPF9 cell association by yeast agglutination assay. Both UTI89 and KqPF9 were able to agglutinate *Saccharomyces cerevisiae* strain L40. However, while treatment with D-mannose abrogated agglutination of yeast by UTI89, we observed no effect of D-mannose on KqPF9-mediated yeast agglutination (Figure S1) (43). Together, these results suggest that KqPF9 association and invasion of bladder epithelial cells and yeast agglutination is mannose insensitive and therefore may not rely on type 1 fimbriae.

**Figure 1.**
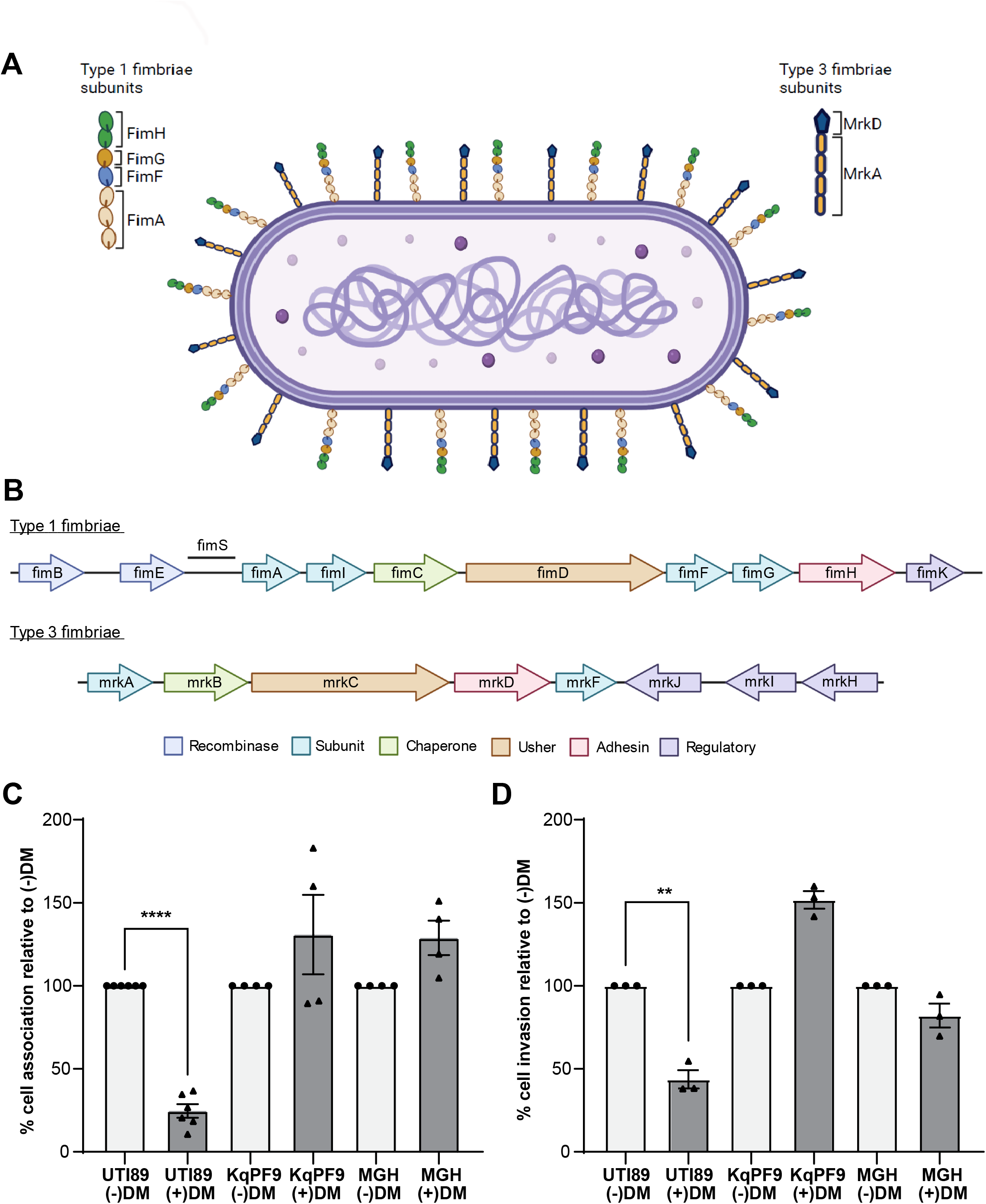
*K. quasipneumoniae* association and invasion to bladder epithelial cells is mannose insensitive. **A**) Illustration of *Klebsiella* type 1 and type 3 fimbriae. FimA and FimH, MrkA and MrkD represent the major structural subunit and adhesin for type1 and 3 fimbriae, respectively. FimF and FimG represent minor structural subunits of type 1 fimbriae. **B**) Schematic of the *fim* and *mrk* operons encoding for type 1 and type 3 fimbriae of *K. quasipneumoniae* isolate PF9 (KqPF9) respectively. Illustrations were made using Biorender.com. **C, D)** 5637 bladder epithelial cell association and invasion in the absence (-DM) and presence (+DM) of 2.5% D-mannose. Uropathogenic *E. coli* strain UTI89 was used as mannose-sensitive control and *K. pneumoniae* type strain MGH78578 was used as mannose-insensitive control. Bladder epithelial cell association of PBS control was evaluated as 100% and the effect of mannose was determined relative to respective PBS control. Experiments were performed in biological and technical triplicate. The error bars indicate standard error of mean. Significance was evaluated by two-tailed, paired Students T test. ** denotes *p<0*.*01* and **** denotes *p<0*.*0005*.

### Type 3 but not type 1 fimbriae are required for *K. quasipneumoniae* attachment to cultured bladder epithelial cells

To evaluate the role of type 1 and type 3 fimbriae of KqPF9 in bladder epithelial cell association, we generated strains with targeted deletions of *fimA* and *mrkA*, which encode for the major fimbrial subunits of the type 1 and 3 fimbrial operons, respectively (34, 44-46). Gene deletions were confirmed by PCR and target gene expression was measured in each mutant and complement strain by qRT-PCR (Figure S2). We next used negative stain electron microscopy to evaluate fimbrial structures produced by each mutant. While fimbrial structures were apparent in wild-type KqPF9 and isogenic type 1 fimbriae mutant KqPF9*ΔfimA*, no fimbrial structures were visible in the isogenic type 3 fimbriae mutant KqPF9*ΔmrkA* or double mutant strain KqPF9*ΔmrkAΔfimA* (Figure 2A). These data suggest that KqPF9 may be preferentially expressing type 3 fimbriae during static growth in LB. Further, fimbrial structures were visible in all complemented mutant strains (Fig 2A). We then performed cell association assays with cultured 5637 bladder epithelial cells to determine the contribution of type 1 and type 3 fimbriae to KqPF9 cell adhesion. The isogenic type 1 fimbriae mutant KqPF9*ΔfimA* strain showed no significant alteration in bladder epithelial cell association relative to wild-type (Figure 2B). However, as previously reported in *K. pneumoniae*, complementation by overexpression of type 1 fimbrial gene cluster (*fimAICDFGHK)* significantly increased bladder epithelial cell association (19). A significant 92.4% decrease in cell association was observed in the double fimbriae mutant KqPF9*ΔmrkAΔfimA*, suggesting that type 3 fimbriae may contribute to bladder epithelial cell association *in vitro* (Figure 2B). Complementation with type 1 fimbrial gene cluster (*fimAICDFGHK)* was able to rescue the cell association phenotype in KqPF9*ΔmrkAΔfimA* strain, which can be attributed to overexpression of the operon (Figure S2D). In contrast to KqPF9*ΔfimA*, the type 3 fimbriae mutant KqPF9*ΔmrkA* showed a significant 71.6% reduction in bladder epithelial cell association, which was complemented by overexpression of the type 3 fimbrial gene cluster (*mrkABCDF*) (Figure 2C). The reduction of cell association in KqPF9*ΔmrkA* and the KqPF9*ΔmrkAΔfimA* double fimbriae mutant was also rescued by overexpression of the *mrkABCDF* gene cluster (Fig 2C). Taken together, these data suggest that *in vitro* association of KqPF9 with bladder epithelial cells is dependent on type 3 but not type 1 fimbriae.

**Figure 2.**
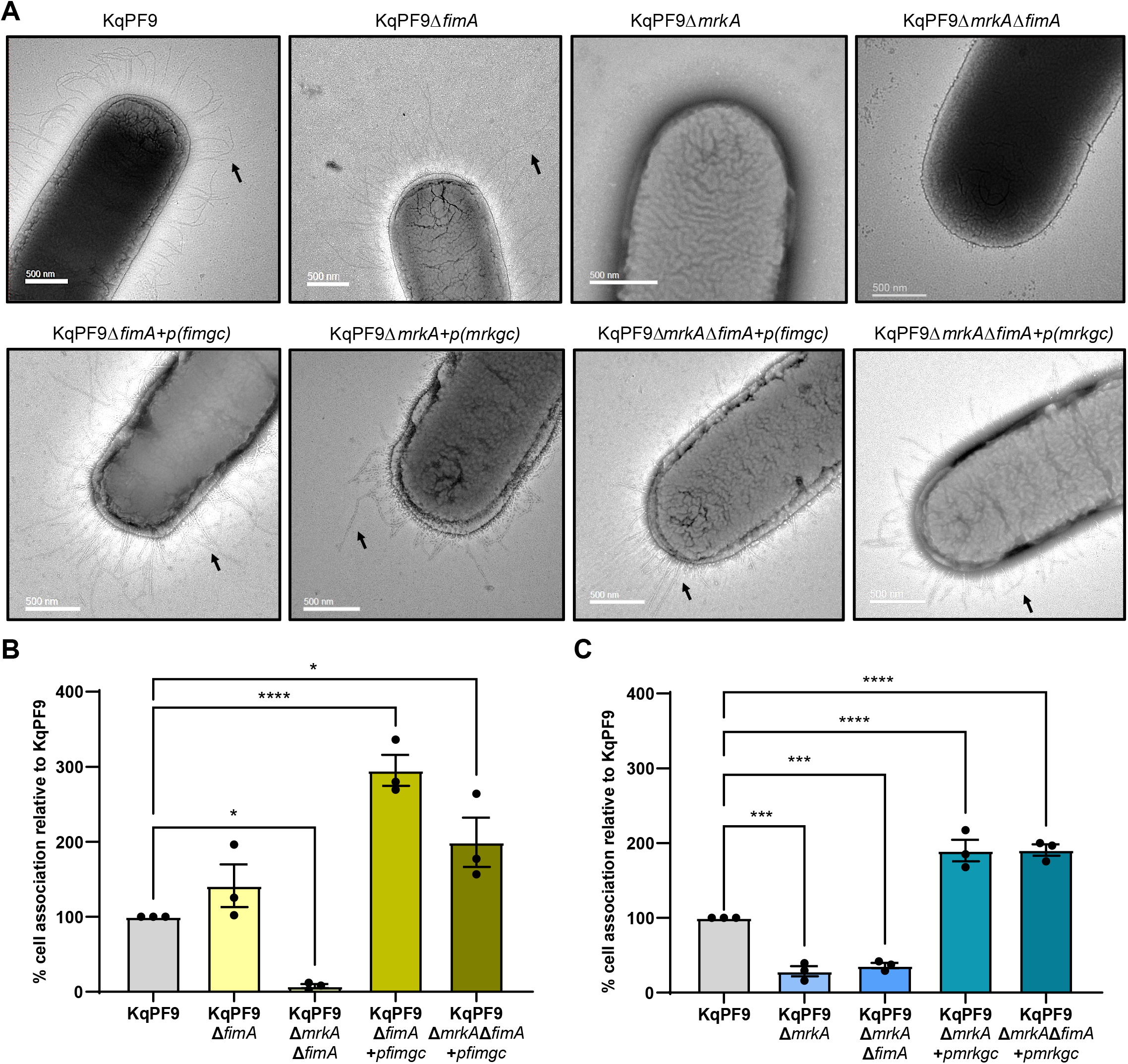
*K. quasipneumoniae* association to cultured bladder epithelial cells is dependent on type 3 fimbriae. **A)** Representative electron micrographs of KqPF9 and isogenic KqPF9Δ*fimA*, KqPF9Δ*mrkA*, and KqPF9Δ*fimA*Δ*mrkA* mutant and complement strains. *fimgc* indicates the *fim* gene cluster (*fimAICDFGHK*) and *mrkgc* indicates the *mrk* gene cluster (*mrkABCDF*). Arrows point to fimbriae. Scale bar is 500nm. **B, C)** Cell association of KqPF9, isogenic type 1 and type 3 fimbriae mutants and mutants expressing respective complementing plasmids to 5637 bladder epithelial cells. The cell association of respective mutants were evaluated relative to KqPF9. The experiments were performed in biological and technical triplicate. The error bars indicate standard error of mean. Significance was evaluated using one way ANOVA with Dunnett’s multiple comparisons post hoc. \**p<0*.*05*, \*\*\**p<0*.*005* and \*\*\*\**p<0*.*0005*.

### Elevated fimbrial expression and bladder epithelial cell association in KqPF9 compared to *K. pneumoniae* TOP52

Because variation in urovirulence phenotypes has been previously reported in *K. variicola*, which was also recently speciated from *K. pneumoniae*, we next wanted to directly compare fimbrial expression and bladder epithelial cell association between KqPF9 and the well-studied uropathogenic *K. pneumoniae* strain TOP52 (2, 27). Quantitative RT-PCR analysis of *fimA* and *mrkA* expression in statically cultured TOP52 and KqPF9 showed a 1.5-fold higher *fimA* expression and 7.4 fold higher *mrkA* expression in KqPF9 relative to TOP52 (Figure 3A, 3B). We then compared cell association frequencies between the two strains and observed that relative bladder epithelial association of KqPF9 was 324.1% higher than TOP52 (Figure 3C). Because we observed significantly increased expression of the major type 3 fimbrial subunit *mrkA* in KqPF9, and biofilm formation has been previously attributed to type 3 fimbriae in *K. pneumoniae*, we also evaluated biofilm formation in each strain (47). In accordance with the elevated *mrkA* expression in KqPF9, we also observed a 744% increase in biofilm formation in KqPF9 with respect to TOP52 (Figure 3D). Because expression of the *fim* operon is controlled by inversion of the *fimS* regulatory element, we performed a *fimS* phase assay and observed that statically grown TOP52 and KqPF9 contained populations both in the “on” and “off” orientation (Figure S3).

**Figure 3.**
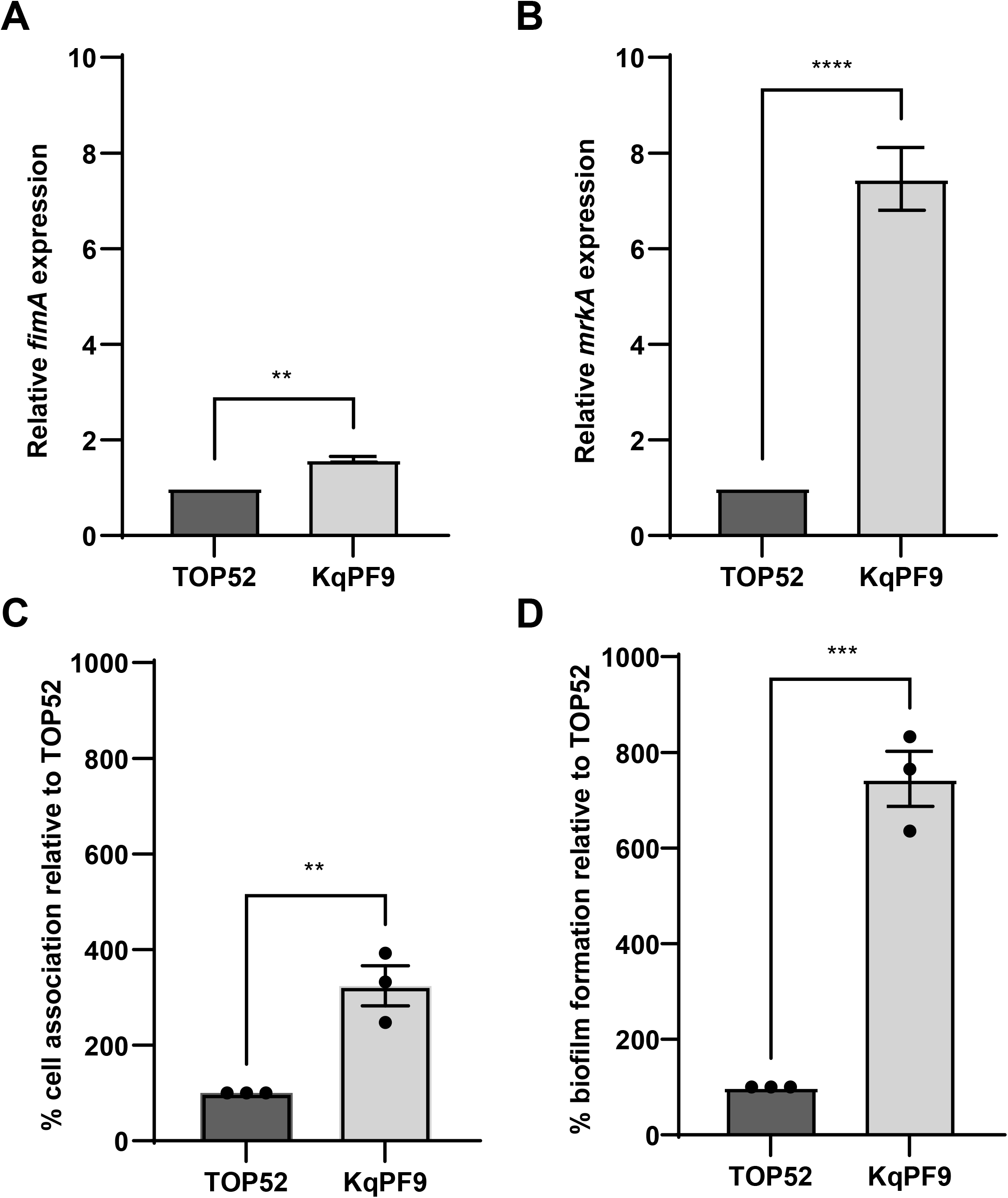
Expression of type 1 and type 3 fimbriae as well as bladder epithelial cell attachment and biofilm formation is elevated in KqPF9 relative to TOP52. **A, B)** qRT-PCR analysis of type 1 fimbriae (*fimA*) and type 3 fimbriae (*mrkA*) expression in TOP52 and KqPF9 isolates. The expression values of *fimA* and *mrkA* were normalized to *rho* and the fold change was determined relative to TOP52. **C)** Percent association KqPF9 with 5637 bladder epithelial cells relative to TOP52. **3D**. Static biofilm formation of TOP52 and KqPF9. Percent KqPF9 biofilm formation was determined relative to TOP52. All experiments were performed at least in biological and technical triplicate. All error bars indicate standard error of mean and two-tailed, paired Student’s T test was used to evaluate significance. \*\**p<0*.*01*, \*\*\**p<0*.*005* and \*\*\*\**p<0*.*0005*.

### The type 1 fimbrial regulatory gene *fimK* is truncated in *K. quasipneumoniae*

We next used previously generated whole genome sequences of TOP52 and KqPF9 to identify genotypic differences explaining the increased expression of *mrkA* and bladder epithelial cell association observed in KqPF9 (5, 48, 49). While we observed no differences in *mrk* operon structure between the two *Klebsiella* species, we found that the type 1 fimbrial regulatory gene, *fimK*, was truncated in KqPF9 (Figure 4A). Indeed, multiple sequence alignment of the FimK amino acid sequences extracted from 10 *K. pneumoniae* and 10 *K. quasipneumoniae* genomes deposited in the NCBI database revealed that the premature stop codon observed in KqPF9 was conserved among *K. quasipneumoniae* isolates resulting in a 218aa protein as compared to the 470aa protein observed in all *K. pneumoniae* isolates (Figure 4B,C) (50, 51). The *fimK* gene of *K. pneumoniae* encodes an N-terminal helix-turn-helix (HTH) DNA binding domain and a C-terminal phosphodiesterase (PDE) domain (39). The premature stop codon observed in *K. quasipneumoniae* results in complete truncation of the C-terminal PDE domain. Because cyclic-di-GMP regulates the expression of type 3 fimbriae through MrkH, it has been hypothesized that the FimK PDE domain may regulate type 3 fimbrial expression by modulating cyclic-di-GMP levels (36, 39). We therefore hypothesized that the elevated *mrkA* expression and bladder epithelial cell association observed in KqPF9 may be due to the absence of the C-terminal PDE domain that is present in *K. pneumoniae* FimK.

**Figure 4.**
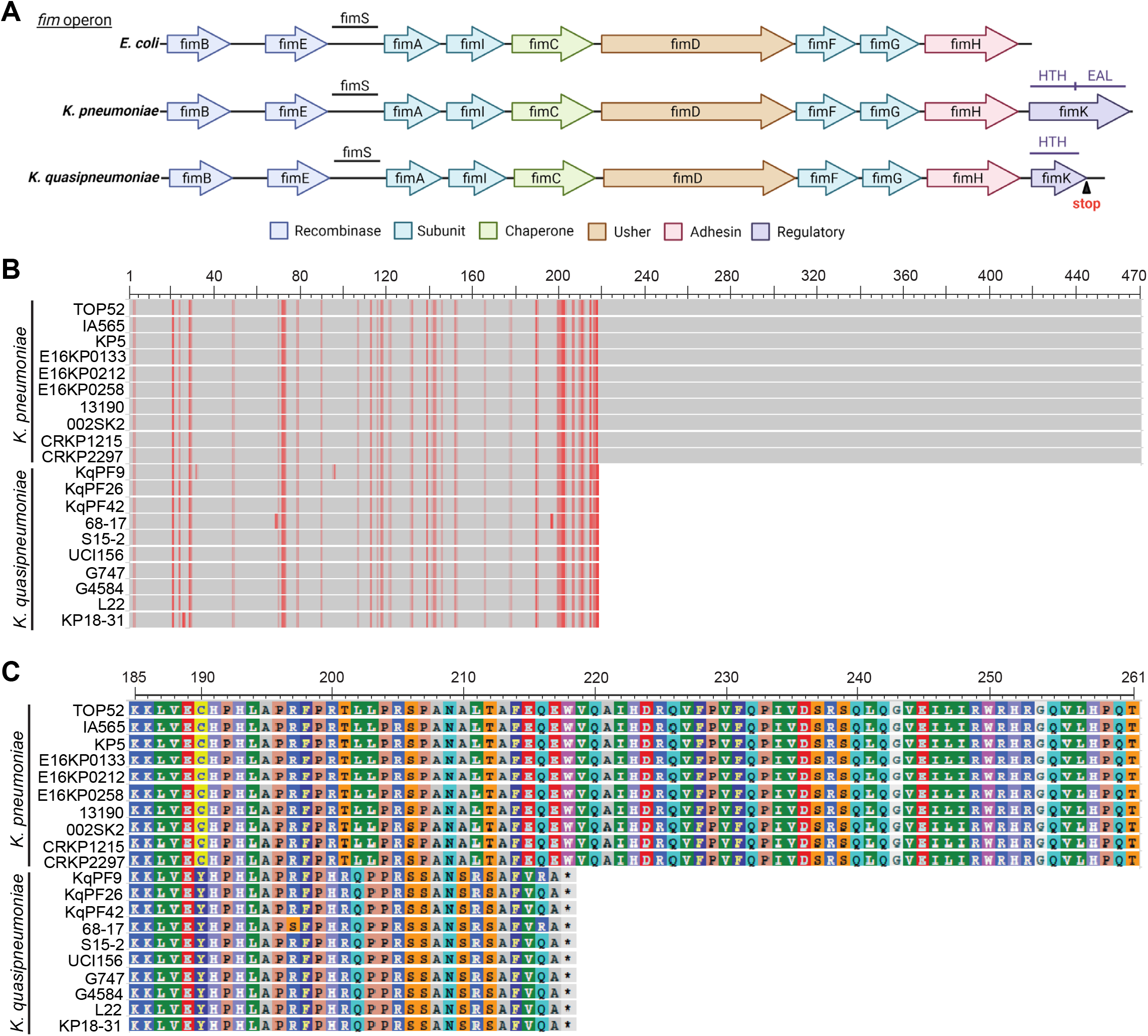
The FimK C-terminal PDE is truncated in *K. quasipneumonaie*. **A)** Comparison of the *fim* operon encoding type 1 fimbriae among the Enterobacteriaceae, *E. coli, K. pneumoniae* and *K. quasipneumoniae. E. coli* does not encode *fimK* and *K. quasipneumoniae* encodes a truncated *fimK* encompassing only the putative helix-turn-helix (HTH) domain with a premature stop codon truncating the EAL phosphodiesterase domain. Schematic made using Biorender.com. **B)** Amino acid alignment of FimK sequences of *K. pneumoniae* and *K. quasipneumoniae* strains deposited in NCBI database is shown with respective sequence IDs. The start and end positions indicate the beginning and end of the FimK amino acid sequence. Mismatches are indicated in red. **C)** A magnification of the alignment in B encompassing the region between amino acids 185 – 261 showing the truncation of *K. quasipneumoniae* FimK at position 218.

### Role of *mrkH* and *mrkJ* in *K. quasipneumoniae* type 3 fimbriae expression

Before investigating differences in FimK function between KqPF9 and TOP52, we first sought to determine if known regulators of type 3 fimbrial expression, MrkH and MrkJ, functioned similarly in *K. quasipneumoniae* as in *K. pneumoniae*. MrkH and MrkJ function respectively as the major activator and repressor of the *mrkABCDF* gene cluster in *K. pneumoniae* (Figure 5A) (35, 36). To test the functions of MrkH and MrkJ in *K. quasipneumoniae*, we generated isogenic KqPF9*ΔmrkH* and KqPF9*ΔmrkJ* mutant strains (Figure S4). We also measured *mrkH* and *mrkJ* expression in mutant and complement strains by qRT-PCR (Figure 5B, 5C). To evaluate the activation and repression of *mrk* gene cluster expression by *mrkH* and *mrkJ* respectively, we studied the expression of *mrkA*. In corroboration with observations previously reported in *K. pneumoniae, mrkA* expression was significantly decreased in the KqPF9*ΔmrkH* strain (2.9-fold) and increased in the KqPF9*ΔmrkJ* strain (2.5-fold) (Figure 5D) (36). In both cases, the respective complement strains, namely KqPF9*ΔmrkH*+*pmrkH* and KqPF9*ΔmrkJ*+*pmrkJ*, rescued *mrkA* expression levels (Figure 5D). Phenotypic analysis showed that bladder epithelial cell association followed a pattern analogous to *mrkA* expression. We observed that the relative bladder epithelial cell association of KqPF9*ΔmrkH* decreased to 2.4% of that observed in wild-type KqPF9 and was rescued upon complementation in KqPF9*ΔmrkH*+*pmrkH* (Figure 5E). Conversely, relative bladder epithelial cell association increased to 217% in KqPF9*ΔmrkJ* compared to wild-type KqPF9. Similar to *mrkA* expression, relative cell association was reduced to 19.1% in the KqPF9*ΔmrkJ*+*pmrkJ* complemented strain compared to wild-type, which was likely due to *mrkJ* overexpression (Figure 5E). Because regulation of type 3 fimbriae by MrkH and MrkJ has been shown to play an important role in biofilm formation in *K. pneumoniae*, we also assessed the role of these regulators in *K. quasipneumoniae* biofilm formation (36). The biofilm assay results followed a similar pattern as cell association (Figure 5F). Relative biofilm formation was significantly decreased in KqPF9*ΔmrkH* to 20.6% and increased to 161% in KqPF9*ΔmrkJ*. These results mirror previously reported findings in uropathogenic *K. pneumoniae* strain AJ218 (36). Taken together, these results suggest that MrkH and MrkJ have similar functions in *K. quasipneumoniae* and *K. pneumoniae* with MrkH acting as an activator of type 3 fimbriae expression and MrkJ acting as a repressor.

**Figure 5.**
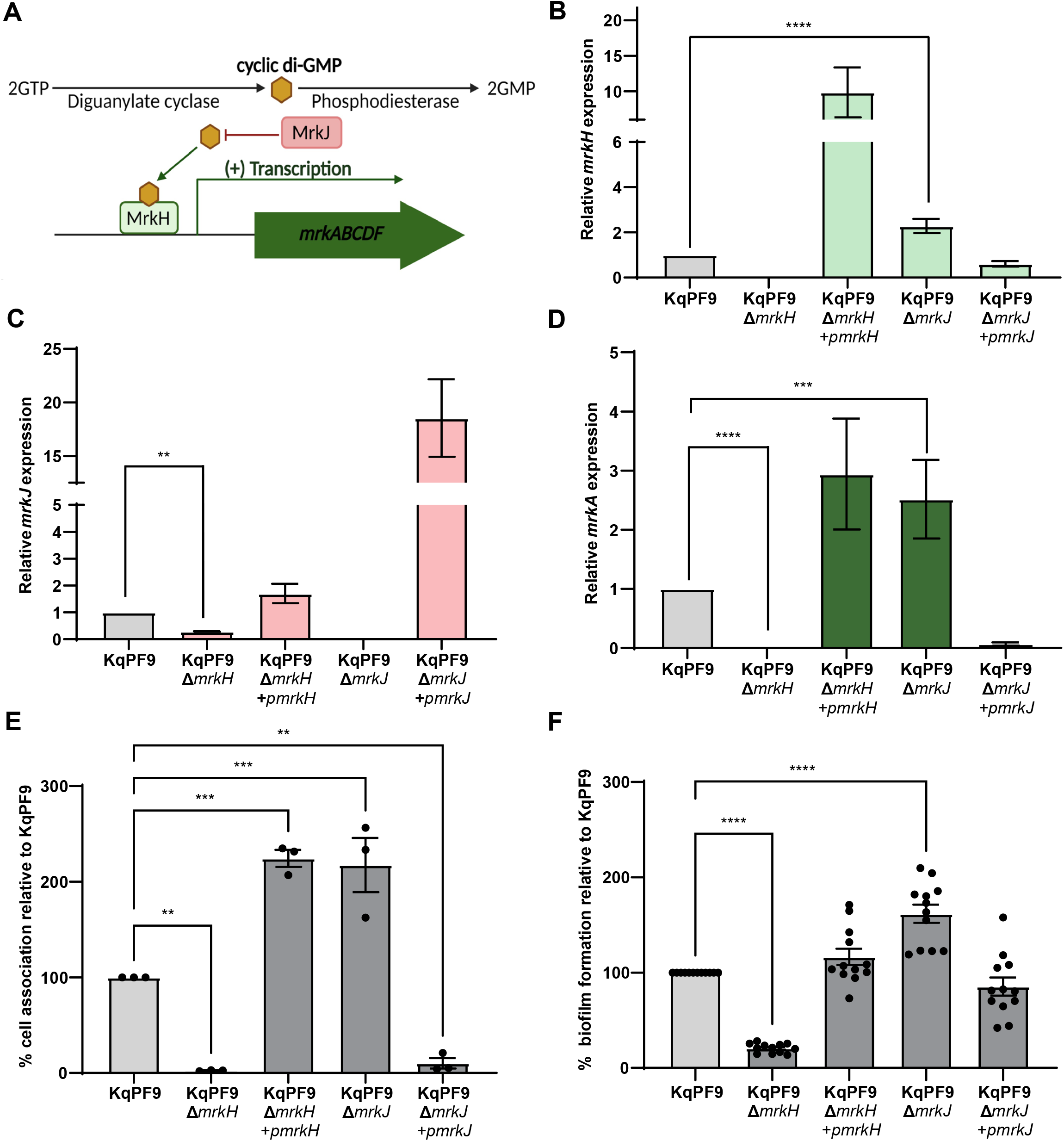
The role of *mrkH* and *mrkJ* in type 3 fimbriae expression is conserved in KqPF9. **A)** Schematic of regulation of the *mrk* operon by transcriptional regulator MrkH, phosphodiesterase MrkJ and cyclic di-GMP in *K. pneumoniae*. qRT-PCR analysis of **B)** *mrkH*, **C)** *mrkJ* and **D)** type 3 fimbriae (*mrkA*) expression in KqPF9Δ*mrkH* and Δ*mrkJ* mutant and complement strains. Expression of each gene was normalized to *rho* and the fold change is expressed relative to wild-type. **E)** Cell association of KqPF9Δ*mrkH* and Δ*mrkJ* mutant and complemented strains to 5637 bladder epithelial cells. The cell association of respective mutant strains was evaluated relative to wild-type KqPF9. **F)** Static biofilm formation of KqPF9 along with isogenic mutants of respective genes, and strains expressing appropriate complementing plasmids. The biofilm formation of the mutant strains was determined relative to KqPF9. All experiments were performed in at least in biological and technical triplicate. The error bars indicate standard error of mean. Statistical testing was performed by one way ANOVA analysis with Dunnett’s multiple comparison test. \*\**p<0*.*01*, \*\*\**p<0*.*005* and \*\*\*\**p<0*.*0005*.

### Overexpression of TOP52 but not KqPF9 *fimK* downregulates expression of type 3 fimbriae

To evaluate the role of the C-terminal PDE domain of FimK in the regulation of type 3 fimbriae expression, we generated TOP52*ΔfimK* strains complemented with plasmids *pfimKKp, pfimK_E245A_* or *pfimK*_*Kq*_ with *fimKKp* encoding the full length TOP52 *fimK* allele, *fimKE245A* encoding a PDE dead allele of TOP52 *fimK*, and *fimK*_*Kq*_ encoding KqPF9 *fimK* allele which lacks the C-terminal PDE domain (Figure S5) (39, 52, 53). Though no significant difference in *mrkH* or *mrkA* expression was observed in between TOP52*ΔfimK* and wild-type TOP52, complementation of TOP52*ΔfimK* with TOP52 *fimK* (*pfimKKp)* showed a significant reduction in *mrkH* (2.94 fold) and *mrkA* (6.25 fold) expression (Figure 6A, 6B). Conversely, complementation of TOP52*ΔfimK* with *pfimK*_*Kq*_ or *pfimK*_*E245A*_, resulted in *mrkH* and *mrkA* expression that remained at levels comparable to wild-type TOP52 (Figure 6A, 6B). Analogous to the changes in *mrkH* and *mrkA* expression, relative bladder epithelial cell association was significantly decreased to 46.14% in TOP52*ΔfimK+pfimKKp* compared to wild-type TOP52 while TOP52*ΔfimK+pfimK_Kq_* and TOP52*ΔfimK+pfimK_E245A_* showed no significant difference in cell association (Figure 6C).

**Figure 6.**
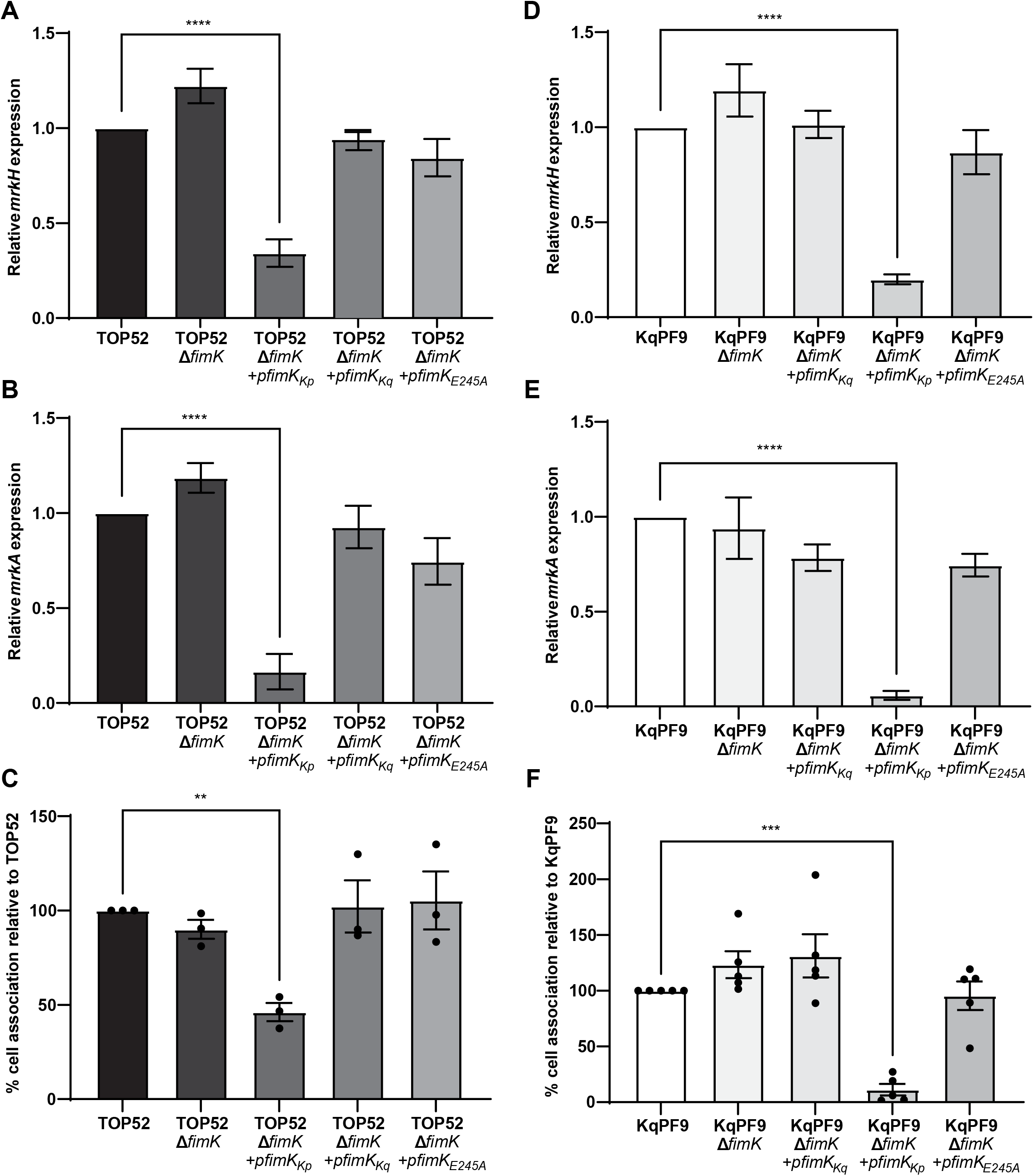
TOP52 *fimK* but not KpPF9 *fimK* reduces type 3 fimbrial expression and bladder epithelial cell attachment. **A, B)**. qRT-PCR analysis of *mrkH* and type 3 fimbriae (*mrkA*) gene transcription in wild-type TOP52 and respective TOP52Δ*fimK* mutant and complemented strains. The expression of each gene was normalized to *rho* and the fold change is relative to wild-type. **C)** 5637 bladder epithelial cell association of TOP52 and respective TOP52Δ*fimK* and complemented strains. **D**,**E)** qRT-PCR evaluating the expression of *mrkH* and *mrkA* in wild-type KqPF9 and respective KqPF9Δ*fimK* mutant and complement strains. *rho* was used for normalization and expression is relative to wild-type. **F)** 5637 bladder epithelial cell association of KqPF9 and KqPF9Δ*fimK* mutant and complemented strains. All experiments were performed at least in biological and technical triplicate. The error bars indicate standard error of mean. One way ANOVA with Dunnett’s multiple comparisons test was used to evaluate significance. \*\**p<0*.*01*, \*\*\**p<0*.*005* and \*\*\*\**p<0*.*0005*.

Considering that only expression of the *fimK* allele containing an intact C-terminal PDE reduced type 3 fimbriae expression in TOP52, we hypothesized that its absence in KqPF9 FimK would translate to an inability to regulate type 3 fimbrial expression in *K. quasipneumoniae*. We generated an isogenic *ΔfimK* mutant of KqPF9 along with complement strains containing either plasmids *pfimK*_*Kq*_, *pfimKKp* or *pfimK*_*E245A*_. Similar to TOP52, significant decreases in *mrkH* (5.2 fold) and *mrkA* (20 fold) expression were observed in KqPF9*ΔfimK+pfimKKp* compared to wild-type KqPF9 (Figure 6D, 6E). However, complementation of KqPF9*ΔfimK* with the PDE-dead TOP52 *fimK* allele (*fimKE245A)* or KqPF9 *fimK* did not alter *mrkA* or *mrkH* expression from wild-type levels (Figure 6D, 6E). Similarly, relative bladder epithelial cell association decreased to 11.2% in KqPF9*ΔfimK+pfimKKp* whereas no change from wild-type KqPF9 was observed in the association of KqPF9*ΔfimK+pfimK_Kq_* and KqPF9*ΔfimK+pfimK_E245A_* (Figure 6F). Taken together these results suggest that while the C-terminal PDE of FimK may function to co-regulate type 3 fimbriae expression in TOP52, truncation of this domain in KqPF9 suggests that this activity is not conserved in *K. quasipneumoniae*.

## Discussion

*K. quasipneumoniae*, previously designated as *K. pneumoniae* phylogroup KpII, was recently classified as a distinct species of *Klebsiella* (37). Previous reports indicate misidentification of *K. quasipneumoniae* as *K. pneumoniae* due to lack of appropriate molecular biology tools in hospital settings (54). Considering that *K. quasipneumoniae* exhibits unique metabolic phenotypes, frequently harbors antibiotic resistance genes like ESBLs and carbapenemases, and is commonly isolated from patients with UTI, a more detailed understanding of the virulence mechanisms involved in bladder colonization of *K. quasipneumoniae* is needed (2).

Both type 1 and type 3 fimbriae have been reported to mediate host cell attachment and invasion by *K. pneumoniae* and are therefore crucial to infection (44, 55). However, no study had measured the expression of type 1 and type 3 fimbriae in *K. quasipneumoniae* and evaluated their respective contributions to bladder epithelial cell attachment. Using isogenic mutants lacking the major subunits of type 1 and type 3 fimbriae we show that *K. quasipneumoniae* strain KqPF9 attachment to cultured bladder epithelial cells is dependent on type 3 but not type 1 fimbriae. While bladder epithelial cell attachment was sharply reduced in KqPF9*ΔmrkA*, no significant difference was observed between KqPF9*ΔfimA* and wild-type KqPF9. Previous reports indicate that type 1, but not type 3 fimbriae are important for *K. pneumoniae* bladder colonization in mice although type 3 fimbriae were found to contribute to urinary bladder epithelial cell attachment *in vitro* (19, 22). It has yet to be determined if type 3 fimbriae are required for *K. quasipneumoniae* bladder colonization in mice – an important subject for future studies.

The regulatory mechanisms controlling the expression of type 1 and type 3 fimbriae in *Klebsiella spp*. are likely more complex than our current understanding. Intracellular second messengers like cyclic-di-GMP play a crucial role in responding to sensory input from extracellular stimuli and coupling them to physiologic changes (56-58). Diguanylate cyclases coordinate the synthesis of cyclic-di-GMP while PDEs with conserved EAL domains are responsible for cyclic-di-GMP degradation (59-62). The type 3 fimbriae expression in *Klebsiella spp*. is to a large extent dependent on transcriptional activator MrkH, a cyclic-di-GMP binding protein, and the MrkJ PDE (36). Interestingly, the FimK protein of *K. pneumoniae* has two domains, an N-terminal DNA binding domain and a C-terminal PDE domain that has been shown to hydrolyze cyclic-di-GMP and thereby regulate cyclic-di-GMP levels (39). It is therefore possible that *K. pneumoniae* FimK contributes to the regulation of type 3 fimbriae by modulating cyclic-di-GMP levels.

The dependence of *K. quasipneumoniae* bladder epithelial cell association on type 3 fimbriae could be attributed to differences in expression due to differential regulation of type 1 and type 3 fimbriae between the two species. Indeed, we observed that basal expression of *mrkA* as well as rates of bladder epithelial cell association were significantly higher in KqPF9 than in *K. pneumoniae* TOP52. Through comparative genomics of the type 1 fimbrial operons of several *K. pneumoniae* and *K. quasipneumoniae* strains, we discovered a premature stop codon in KqPF9 *fimK* that results in truncation of the C-terminal PDE domain and is conserved among *K. quasipneumoniae* isolates. We further demonstrated the role of the TOP52 FimK PDE in regulating type 3 fimbriae expression and bladder epithelial cell attachment *in vitro*. These results suggest that the higher basal expression of type 3 fimbriae in KqPF9 relative to TOP52 may at least be, in part, attributed to FimK PDE domain truncation. We also show that MrkH and MrkJ have conserved respective activator and repressor functions in the regulation of type 3 fimbriae in *K. quasipneumoniae*.

Interestingly, we found that deletion of *fimK* did not significantly elevate *mrkA* expression in TOP52. We hypothesize that this may be due to functional redundancy with other phosphodiesterases like MrkJ. However, it is possible that the presence of other diguanylate cyclases and phosphodiesterases in *Klebsiella quasipneumoniae* may contribute to co-regulation of type 1 and type 3 fimbriae in different host environments. Also, previous work has shown that FimK inhibits type 1 fimbrial expression in TOP52 and that TOP52*ΔfimK* exhibits significantly increased type 1 fimbrial expression and bladder colonization (27). The absence of PDE domain in *fimK* of KqPF9 alongside an increase in expression of type 1 (*fimA*) and type 3 (*mrkA*) fimbriae may explain the increased biofilm formation and bladder epithelial cell association of this strain relative to TOP52. However, it is yet unclear if the increased cell association of KqPF9 strain translates to increased colonization of the urinary tract *in vivo* and if KqPF9 bladder colonization is dependent on type 1 or type 3 fimbriae.

It is also possible that additional regulatory differences exist between the two species that may contribute to differences in expression of type 1 and type 3 fimbriae. In addition to local regulators of type 1 and type 3 fimbriae encoded within the *fim* and *mrk* operons, global regulators like integration host factor (IHF), leucine regulatory proteins (LRP) and histone-like nucleoid structuring protein (H-NS) also play an important role in fimbrial regulation (63-66). In *E. coli*, IHF and LRP are required to maintain the phase on orientational bias of the *fimS* switch controlling type 1 fimbriae expression while H-NS is required to maintain the phase off orientation (66). H-NS has been reported to regulate type 1 fimbriae in *E. coli* and *K. pneumoniae* and type 3 fimbriae in *K. pneumoniae* (63, 64, 67). H-NS regulates the phase variation of the type 1 fimbriae *fimS* region directly by binding to region adjacent to *fimS* (68-70). H-NS also regulates type 1 fimbriae indirectly by repressing the transcription of *fimB* and *fimE* recombinases (71). In case of type 3 fimbriae, H-NS represses the expression of *mrkH* and *mrkI* (63) and activates *mrkA* expression (72). Interestingly, cyclic-di-GMP has been reported to alleviate H-NS repression of genes involved in biofilm formation in *Vibrio* cholerae (73). Considering that the MrkH activator is dependent on cyclic-di-GMP, it is possible that cyclic-di-GMP activates regulators like MrkH whose binding site overlaps with binding sites of H-NS (72, 74) thereby coordinating de-repression of genes involved in specific function that were originally repressed by H-NS (47, 72). Future studies should focus on disentangling these additional regulatory mechanisms in *K. quasipneumoniae*.

Taken together, the data presented here provide new insight into the regulation of type 3 fimbriae by *fimK* in *K. pneumoniae* as well as provides evidence for important differences in fimbrial expression and bladder epithelial cell attachment phenotypes between *K. pneumoniae* and *K. quasipneumoniae* species.

## Materials and Methods

### Bacterial strains and culture conditions

Bacterial strains and plasmids used in the study are listed in Table 1. KqPF9, a *K. quasipneumoniae* strain isolated from the urine of a postmenopausal woman with active, symptomatic rUTI (5), TOP52, a *K. pneumoniae* strain isolated from urine of a woman with acute cystitis and UTI89 an UPEC acute cystitis isolate were used in the study. The complete KqPF9 genome has been described previously and is available on GenBank through BioProject PRJNA683049 (5). Bacteria were grown in Luria Bertani (LB) media overnight at 37°C in static conditions. For ribonucleic acid (RNA) extractions in quantitative reverse transcriptase polymerase chain reaction (qRT-PCR) assays, overnight cultures were sub-cultured at a starting density of OD_600_ = 0.01 and then incubated for 6hrs until reaching mid-log phase (OD_600_ = 0.4 – 0.6) induced with 0.2% L-arabinose (*mrkA* genotypes) or 0.0125% L-arabinose (*fimA, mrkH, mrkJ* genotypes) at 37°C in LB media in an incubator at static conditions. For negative staining electron microscopy experiments, bacteria were grown for 8 hrs at 37°C in static conditions with 0.2% L-arabinose for induction of complementation in respective strains.

**Table 1.** Strains and Plasmids used in this study.

| Strain and plasmids | Description | Source |
| --- | --- | --- |
| PF9 | *K. quasipneumoniae* cystitis isolate | This study |
| TOP52 1721 | *K. pneumoniae* cystitis isolate | Rosen et al, 2008 (27) |
| UTI89 | *E. coli* cystitis isolate | Mulvey et al, 2001 (84) |
| KqPF9Δ*mrkA* | Knockout of *mrkA* in PF9 | This study |
| KqPF9Δ*fimA* | Knockout of *fimA* in PF9 | This study |
| KqPF9Δ*mrkA*Δ*fimA* | Knockout of *mrkA* and *fimA* in PF9 | This study |
| KqPF9Δ*mrkH* | Knockout of *mrkH* in PF9 | This study |
| KqPF9Δ*mrkJ* | Knockout of *mrkJ* in PF9 | This study |
| KqPF9Δ*fimK* | Knockout of *fimK* in PF9 | This study |
| TOP52Δ*fimK* | Knockout of *fimK* in TOP52 | This study |
| *pBAD* - empty vector | Empty vector (EV), arabinose inducible, Kan<sup>R</sup> | Guzman et al, 1995 (77) |
| *pmrkgc* | Plasmid expressing *mrk* gene cluster | This study |
| *pfimgc* | Plasmid expressing *fim* gene cluster | This study |
| *pmrkH* | Plasmid expressing *mrkH* | This study |
| *pmrkJ* | Plasmid expressing *mrkJ* | This study |
| *pfimK<sub>Kp</sub>* | Plasmid expressing WT TOP52*fimK* | This study |
| *pfimK<sub>Kq</sub>* | Plasmid expressing WT KqPF9 *fimK* | This study |
| *pfimK<sub>E245A</sub>* | Plasmid expressing mutant TOP52 *fimK*-*AIL* | This study |

### Bacterial cell association and gentamicin protection (invasion) assays

Cell association and invasion assays were performed as previously described (41). Urinary bladder epithelial 5637 cells (ATCC#HTB-9) were cultivated in RPMI (Sigma) supplemented with 10% FBS (ThermoFisher), 1% penicillin-streptomycin (Gibco) and 1% glutamax (Gibco) overnight at 37 °C and 5% CO2. The next day, cells were washed with phosphate buffered saline (PBS) and incubated in RPMI without antibiotics. Cells were infected at a multiplicity of infection (MOI) of 10. Bacterial contact with host cells was synchronized by centrifugation of plates at 600 x g for 5 min. For association assays, 2 hours post-infection cells were washed in 1X PBS and then harvested and lysed with 0.3% Triton X-100. Cells in the input wells, representing total intra and extracellular bacteria were not washed and directly lysed with Triton X-100. For gentamicin protection (invasion) assays, RPMI was replaced with RPMI containing 100 µg/mL gentamicin 2 hours-post infection and incubated for an additional 2 hours before harvesting as described above. Association and invasion frequencies were calculated as a ratio of the number of bacteria recovered from washed lysates to the total number of bacteria present in the input (unwashed) wells. All cell association and invasion assays were performed at least in biological triplicate. For each biological replicate, a minimum of 3 technical replicates were evaluated.

### Generation of *K. quasipneumoniae* and *K. pneumoniae* gene deletion mutants

Type 1 fimbriae knockout, type 3 fimbriae knockout or both type 1 and type 3 fimbriae knockout mutants of *K. quasipneumoniae* were generated by targeted gene deletion of genes encoding the major fimbrial subunits, *fimA* and *mrkA*, respectively (34, 44). All gene deletions were performed using Lambda Red Recombinase system as previously described with respective primers listed in Table S1 (75). The plasmid pKD4 served as the template to amplify the kanamycin cassette and flanking FRT sites (76) to generate knockout cassettes respective to each gene. The Lambda Red Recombinase system was expressed from pACBSR-Hyg which carries *beta, gam*, and *exo* under the control of an arabinose inducible promoter. pFLP-Hyg, which expresses the FLP recombinase, was used to excise the kanamycin cassette via the FRT sites. PCR was performed using flanking primers mentioned in Table S1 to screen candidate colonies using Dreamtaq Master Mix (ThermoFisher). PCR products were analyzed by agarose gel electrophoresis and Sanger sequencing (Genewiz) to confirm gene deletion.

### Generation of complementation plasmids

For construction of complementation plasmids, genes were amplified from genomic DNA by PCR using Dreamtaq Mastermix (ThermoFisher) or PHUsion polymerase for PCR products >2,000bp (NEB) with respective primers as provided in Supplemental Table 1. For all plasmids, NcoI and HindIII restriction sites were used except for plasmids encoding *mrk* and *fim* gene clusters where EcoRI and XhoI sites were used for ligation into pBAD33 (Kan^R^) version A (77). For the generation of the plasmid encoding the AIL mutant of *fimKE245A*, site-directed mutagenesis was performed on *pfimKKp* with the mutagenesis primers listed in Supplemental Table 1 followed by DpnI digestion. All plasmids were sequence verified by Sanger sequencing before transformation into respective isogenic mutant strains.

### Bacterial transformation

Transformation of *Klebsiella* strains KqPF9 and TOP52 was performed by electroporation of respective electrocompetent cells in 0.1 cm gap-width cuvette, 2.5 kV using a micropulser electroporator (Bio-Rad) and selected on LB agar plates with 100µg/mL kanamycin for Lambda Red Recombineering protocols and selection of complementation plasmids (75). Chemical transformation was used during subcloning to introduce complementation plasmid constructs into *E. coli* DH5α (77).

### RNA extraction, cDNA synthesis and qRT-PCR

Total RNA was extracted from respective statically growing *K*qPF9 and TOP52 at mid-log phase OD_600_ = 0.5-0.6) using the Qiagen RNeasy Plus kit by following manufacturer’s instructions. RNA concentration and quality was analyzed by NanoDrop (Thermo). RNA with A260/280 ratios between 2.0-2.20 was used for cDNA synthesis using the Qscript cDNA kit (Quantabio). Following cDNA synthesis, qPCR was performed using PerfeCTa SYBR Green FastMix (Quantabio) and respective primer pairs listed in Table S2. The expression of the gene of interest was normalized to the expression of *rho* (78). Relative expression was determined by the ΔΔCq method (79, 80). qRT-PCR was performed in biological and technical triplicates.

### Static biofilm assays

Static biofilm assays were performed as described previously (81). Bacteria were grown overnight in LB at 37°C under static conditions. Bacteria were normalized to an OD_600_ of 0.1 in LB broth and seeded into round bottom 96 well plates (Corning). 0.2% L-arabinose was added for induction of complementation plasmids. Plates were incubated for 23 hours at 37°C in static conditions and then washed (2X) with sterile water and dried for 25 minutes. Biofilms were fixed with methanol, stained with a 0.1% (w/v) crystal violet solution (Sigma) and washed (2X) with sterile water. The bound dye was solubilized with 30% acetic acid and the OD_585_ was measured. Each experiment was performed at least three times in technical triplicate.

### Yeast Agglutination Assays

Bacterial strains were grown overnight at 37°C in Luria Bertani (LB) media at static conditions. Cultures were normalized to an OD_600_ of 0.5 and centrifuged at 5,000rpm for 5 minutes. Bacterial pellets were then resuspended in 1X sterile PBS or PBS + 2.5% D-Mannose solutions. *Saccharomyces cerevisiae* strain L40 was grown overnight at 27°C in Yeast Peptone Dextrose (YPD) media at 225 rpm. Yeast cells were normalized to an OD_600_ of 0.5, pelleted at 5,000 rpm and resuspended in in 1X sterile PBS. Bacterial and yeast suspensions were then mixed in a 1:1 ratio on a glass slide and was gently rotated until agglutination became visible and imaged under a light microscope (Zeiss). Images were obtained using XCAM1080PHD and SEBA view software.

### Multiple Sequence Alignments

The *fimK* genes of 10 different *K. pneumoniae* and *K. quasipneumoniae* strains that were previously reported in NCBI database were compiled. The *fimK* sequences of three *K. quasipneumoniae* isolates with sequence IDs CP065841.1 (PF9), CP065838.1 (PF26) and CP065846.1 (PF42) whose whole genome sequencing was performed and previously reported were also used for alignment (5). Following collation of the translated *fimK* sequences, their amino acid sequence alignment was performed using Muscle software with default parameters (50, 51). The alignments were then exported to NCBI’s multiple sequence aligner for further analysis.

### Phase assay of the *fimS* invertible element of *fim* operon

The *fimS* phase assays were performed similarly as described previously (82). KqPF9 and TOP52 isolates were grown overnight at 37°C in static conditions, harvested by centrifugation and their genomic DNA extracted using the Easy-DNA kit (Invitrogen). PCR primers were designed listed in Supplemental Table 1 to amplify nucleotide segments 5’ and 3’ to the respective *fimS* regions. The “on” phase generated a PCR product of size 950bp while the “off” phase generated a product of size 356bp. Based on size of the PCR products as analyzed by 1% agarose gel electrophoresis, the corresponding “on” or “off” phase of the *fimS* switch was determined.

### Negative stain electron microscopy

Negative staining and electron microscopy were performed as previously described (83). Briefly, a formvar-coated carbon-reinforced copper grid was placed with the film side down, on a droplet of a bacterial suspension for 2 minutes. Filter paper was used to remove excess liquid and the grid was stained for 30 seconds on droplets of 1.25% phosphotungstic acid (pH 6.5). Electron microscopy was performed with a JEOL 1400+ transmission electron microscope using Gatan Microscopy Suite (GMS) software.

## Statistical analysis

All statistical analysis was performed using GraphPad Prism (9.1.0). For pairwise comparisons, significance was evaluated by Paired two tailed Student’s ‘T’ test. For multiple comparisons, significance was evaluated using one way ANOVA with Dunnett’s multiple comparison test. For all tests, *p* ≤ 0.05 was considered significant and represented as * while *p* ≤ 0.01 was represented as **, *p* ≤ 0.001 was represented as ***, *p* ≤ 0.0001 was represented as ****.

## Acknowledgements

We would like to sincerely thank David A. Rosen (WUSTL) for providing the *K. pneumoniae* TOP52 strain and Zachary T. Campbell (UTD) for providing *Saccharomyces cerevisiae* strain L40. This work was supported by grants from the Welch Foundation (AT-2030-20200401) and the National Institutes of Health (1R01DK131267-01) to NJD, The Felicia and John Cain Distinguished Chair in Women’s Health to PEZ, as well as grants from the National Science Foundation (DMR-2003534) and the Welch Foundation (AT-1989-20190330) to JJG.

## Author Contributions

S.V. and N.J.D were responsible for conceptualization and experimental design. Primary manuscript writing and editing was done by S.V. and N.J.D. Generation of gene knockouts, plasmid construction, RNA extraction, qRT-PCR, bacterial cell association assay, bacterial invasion assay, static biofilm assay, phase transition assay of *fim* operon, agglutination assays were performed by S.V. Negative stain electron microscopy was performed by Y.H.W. and F.C.H. Statistical analyses and interpretation were performed by S.V. And N.J.D. Funding was raised by N.J.D, P.E.Z. and J.J.G.

## Competing Interest

The authors declare no competing interest.

## Notes

### Competing Interest Statement

The authors have declared no competing interest.

## References

1. Podschun R, Ullmann U. 1998. Klebsiella spp. as nosocomial pathogens: epidemiology, taxonomy, typing methods, and pathogenicity factors. Clin Microbiol Rev 11:589–603.

2. Potter RF, Lainhart W, Twentyman J, Wallace MA, Wang B, Burnham CA, Rosen DA, Dantas G. 2018. Population Structure, Antibiotic Resistance, and Uropathogenicity of Klebsiella variicola. mBio 9.

3. Holt KE, Wertheim H, Zadoks RN, Baker S, Whitehouse CA, Dance D, Jenney A, Connor TR, Hsu LY, Severin J, Brisse S, Cao H, Wilksch J, Gorrie C, Schultz MB, Edwards DJ, Nguyen KV, Nguyen TV, Dao TT, Mensink M, Minh VL, Nhu NT, Schultsz C, Kuntaman K, Newton PN, Moore CE, Strugnell RA, Thomson NR. 2015. Genomic analysis of diversity, population structure, virulence, and antimicrobial resistance in Klebsiella pneumoniae, an urgent threat to public health. Proc Natl Acad Sci U S A 112:E3574–81.

4. Gupta K, Sahm DF, Mayfield D, Stamm WE. 2001. Antimicrobial resistance among uropathogens that cause community-acquired urinary tract infections in women: a nationwide analysis. Clin Infect Dis 33:89–94.

5. Venkitapathi S, Sharon BM, Ratna TA, Arute AP, Zimmern PE, De Nisco NJ. 2021. Complete Genome Sequences of Three Uropathogenic Klebsiella quasipneumoniae Strains Isolated from Postmenopausal Women with Recurrent Urinary Tract Infection. Microbiol Resour Announc 10.

6. Scholes D, Hooton TM, Roberts PL, Stapleton AE, Gupta K, Stamm WE. 2000. Risk factors for recurrent urinary tract infection in young women. J Infect Dis 182:1177–82.

7. Foxman B. 1990. Recurring urinary tract infection: incidence and risk factors. Am J Public Health 80:331–3.

8. Malik RD, Wu YR, Zimmern PE. 2018. Definition of Recurrent Urinary Tract Infections in Women: Which One to Adopt? Female Pelvic Med Reconstr Surg 24:424–429.

9. Cristea OM, Avramescu CS, Balasoiu M, Popescu FD, Popescu F, Amzoiu MO. 2017. Urinary tract infection with Klebsiella pneumoniae in Patients with Chronic Kidney Disease. Curr Health Sci J 43:137–148.

10. Flores-Mireles AL, Walker JN, Caparon M, Hultgren SJ. 2015. Urinary tract infections: epidemiology, mechanisms of infection and treatment options. Nature Reviews Microbiology 13:269–284.

11. Langley JM, Hanakowski M, Leblanc JC. 2001. Unique epidemiology of nosocomial urinary tract infection in children. Am J Infect Control 29:94–8.

12. Marschall J, Fraser VJ, Doherty J, Warren DK. 2009. Between community and hospital: healthcare-associated gram-negative bacteremia among hospitalized patients. Infect Control Hosp Epidemiol 30:1050–6.

13. Paterson DL, Hujer KM, Hujer AM, Yeiser B, Bonomo MD, Rice LB, Bonomo RA, International Klebsiella Study G. 2003. Extended-spectrum beta-lactamases in Klebsiella pneumoniae bloodstream isolates from seven countries: dominance and widespread prevalence of SHV- and CTX-M-type beta-lactamases. Antimicrob Agents Chemother 47:3554–60.

14. Nordmann P, Cuzon G, Naas T. 2009. The real threat of Klebsiella pneumoniae carbapenemase-producing bacteria. Lancet Infect Dis 9:228–36.

15. Partridge SR, Thomas LC, Ginn AN, Wiklendt AM, Kyme P, Iredell JR. 2011. A novel gene cassette, aacA43, in a plasmid-borne class 1 integron. Antimicrob Agents Chemother 55:2979–82.

16. Deguchi T, Fukuoka A, Yasuda M, Nakano M, Ozeki S, Kanematsu E, Nishino Y, Ishihara S, Ban Y, Kawada Y. 1997. Alterations in the GyrA subunit of DNA gyrase and the ParC subunit of topoisomerase IV in quinolone-resistant clinical isolates of Klebsiella pneumoniae. Antimicrob Agents Chemother 41:699–701.

17. Karah N, Poirel L, Bengtsson S, Sundqvist M, Kahlmeter G, Nordmann P, Sundsfjord A, Samuelsen O, Norwegian Study Group on P. 2010. Plasmid-mediated quinolone resistance determinants qnr and aac(6’)-Ib-cr in Escherichia coli and Klebsiella spp. from Norway and Sweden. Diagn Microbiol Infect Dis 66:425–31.

18. Schroll C, Barken KB, Krogfelt KA, Struve C. 2010. Role of type 1 and type 3 fimbriae in Klebsiella pneumoniae biofilm formation. BMC Microbiol 10:179.

19. Struve C, Bojer M, Krogfelt KA. 2009. Identification of a conserved chromosomal region encoding Klebsiella pneumoniae type 1 and type 3 fimbriae and assessment of the role of fimbriae in pathogenicity. Infect Immun 77:5016–24.

20. Krogfelt KA, Bergmans H, Klemm P. 1990. Direct evidence that the FimH protein is the mannose-specific adhesin of Escherichia coli type 1 fimbriae. Infect Immun 58:1995–8.

21. Klemm P, Krogfelt KA, Hedegaard L, Christiansen G. 1990. The major subunit of Escherichia coli type 1 fimbriae is not required for D-mannose-specific adhesion. Mol Microbiol 4:553–9.

22. Tarkkanen AM, Virkola R, Clegg S, Korhonen TK. 1997. Binding of the type 3 fimbriae of Klebsiella pneumoniae to human endothelial and urinary bladder cells. Infect Immun 65:1546–9.

23. Oelschlaeger TA, Tall BD. 1997. Invasion of cultured human epithelial cells by Klebsiella pneumoniae isolated from the urinary tract. Infect Immun 65:2950–8.

24. Duguid JP, Smith IW, Dempster G, Edmunds PN. 1955. Non-flagellar filamentous appendages (fimbriae) and haemagglutinating activity in Bacterium coli. J Pathol Bacteriol 70:335–48.

25. Duguid JP. 1959. Fimbriae and adhesive properties in Klebsiella strains. J Gen Microbiol 21:271–86.

26. Gerlach GF, Clegg S, Ness NJ, Swenson DL, Allen BL, Nichols WA. 1989. Expression of type 1 fimbriae and mannose-sensitive hemagglutinin by recombinant plasmids. Infect Immun 57:764–70.

27. Rosen DA, Pinkner JS, Jones JM, Walker JN, Clegg S, Hultgren SJ. 2008. Utilization of an intracellular bacterial community pathway in Klebsiella pneumoniae urinary tract infection and the effects of FimK on type 1 pilus expression. Infect Immun 76:3337–45.

28. Abraham JM, Freitag CS, Clements JR, Eisenstein BI. 1985. An invertible element of DNA controls phase variation of type 1 fimbriae of Escherichia coli. Proc Natl Acad Sci U S A 82:5724–7.

29. Klemm P. 1986. Two regulatory fim genes, fimB and fimE, control the phase variation of type 1 fimbriae in Escherichia coli. EMBO J 5:1389–93.

30. Adegbola RA, Old DC. 1982. New fimbrial hemagglutinin in Serratia species. Infect Immun 38:306–15.

31. Burmolle M, Bahl MI, Jensen LB, Sorensen SJ, Hansen LH. 2008. Type 3 fimbriae, encoded by the conjugative plasmid pOLA52, enhance biofilm formation and transfer frequencies in Enterobacteriaceae strains. Microbiology (Reading) 154:187–195.

32. Norman A, Hansen LH, She Q, Sorensen SJ. 2008. Nucleotide sequence of pOLA52: a conjugative IncX1 plasmid from Escherichia coli which enables biofilm formation and multidrug efflux. Plasmid 60:59–74.

33. Gerlach GF, Allen BL, Clegg S. 1988. Molecular characterization of the type 3 (MR/K) fimbriae of Klebsiella pneumoniae. J Bacteriol 170:3547–53.

34. Murphy CN, Clegg S. 2012. Klebsiella pneumoniae and type 3 fimbriae: nosocomial infection, regulation and biofilm formation. Future Microbiol 7:991–1002.

35. Johnson JG, Clegg S. 2010. Role of MrkJ, a phosphodiesterase, in type 3 fimbrial expression and biofilm formation in Klebsiella pneumoniae. J Bacteriol 192:3944–50.

36. Wilksch JJ, Yang J, Clements A, Gabbe JL, Short KR, Cao H, Cavaliere R, James CE, Whitchurch CB, Schembri MA, Chuah ML, Liang ZX, Wijburg OL, Jenney AW, Lithgow T, Strugnell RA. 2011. MrkH, a novel c-di-GMP-dependent transcriptional activator, controls Klebsiella pneumoniae biofilm formation by regulating type 3 fimbriae expression. PLoS Pathog 7:e1002204.

37. Brisse S, Passet V, Grimont PAD. 2014. Description of Klebsiella quasipneumoniae sp. nov., isolated from human infections, with two subspecies, Klebsiella quasipneumoniae subsp. quasipneumoniae subsp. nov. and Klebsiella quasipneumoniae subsp. similipneumoniae subsp. nov., and demonstration that Klebsiella singaporensis is a junior heterotypic synonym of Klebsiella variicola. Int J Syst Evol Microbiol 64:3146–3152.

38. Imai K, Ishibashi N, Kodana M, Tarumoto N, Sakai J, Kawamura T, Takeuchi S, Taji Y, Ebihara Y, Ikebuchi K, Murakami T, Maeda T, Mitsutake K, Maesaki S. 2019. Clinical characteristics in blood stream infections caused by Klebsiella pneumoniae, Klebsiella variicola, and Klebsiella quasipneumoniae: a comparative study, Japan, 2014-2017. BMC Infect Dis 19:946.

39. Wang ZC, Huang CJ, Huang YJ, Wu CC, Peng HL. 2013. FimK regulation on the expression of type 1 fimbriae in Klebsiella pneumoniae CG43S3. Microbiology (Reading) 159:1402–1415.

40. Martinez JJ, Mulvey MA, Schilling JD, Pinkner JS, Hultgren SJ. 2000. Type 1 pilus-mediated bacterial invasion of bladder epithelial cells. EMBO J 19:2803–12.

41. Eto DS, Jones TA, Sundsbak JL, Mulvey MA. 2007. Integrin-mediated host cell invasion by type 1-piliated uropathogenic Escherichia coli. PLoS Pathog 3:e100.

42. Rosen DA, Pinkner JS, Walker JN, Elam JS, Jones JM, Hultgren SJ. 2008. Molecular variations in Klebsiella pneumoniae and Escherichia coli FimH affect function and pathogenesis in the urinary tract. Infect Immun 76:3346–56.

43. Stahlhut SG, Struve C, Krogfelt KA. 2012. Klebsiella pneumoniae type 3 fimbriae agglutinate yeast in a mannose-resistant manner. J Med Microbiol 61:317–322.

44. Struve C, Bojer M, Krogfelt KA. 2008. Characterization of Klebsiella pneumoniae type 1 fimbriae by detection of phase variation during colonization and infection and impact on virulence. Infect Immun 76:4055–65.

45. Huynh DTN, Kim AY, Kim YR. 2017. Identification of Pathogenic Factors in Klebsiella pneumoniae Using Impedimetric Sensor Equipped with Biomimetic Surfaces. Sensors (Basel) 17.

46. Wang H, Wilksch JJ, Chen L, Tan JW, Strugnell RA, Gee ML. 2017. Influence of Fimbriae on Bacterial Adhesion and Viscoelasticity and Correlations of the Two Properties with Biofilm Formation. Langmuir 33:100–106.

47. Langstraat J, Bohse M, Clegg S. 2001. Type 3 fimbrial shaft (MrkA) of Klebsiella pneumoniae, but not the fimbrial adhesin (MrkD), facilitates biofilm formation. Infect Immun 69:5805–12.

48. Wylie KM, Wylie TN, Minx PJ, Rosen DA. 2019. Whole-Genome Sequencing of Klebsiella pneumoniae Isolates to Track Strain Progression in a Single Patient With Recurrent Urinary Tract Infection. Front Cell Infect Microbiol 9:14.

49. Johnson JG, Spurbeck RR, Sandhu SK, Matson JS. 2014. Genome Sequence of Klebsiella pneumoniae Urinary Tract Isolate Top52. Genome Announc 2.

50. Edgar RC. 2004. MUSCLE: a multiple sequence alignment method with reduced time and space complexity. BMC Bioinformatics 5:113.

51. Edgar RC. 2004. MUSCLE: multiple sequence alignment with high accuracy and high throughput. Nucleic Acids Res 32:1792–7.

52. Kuchma SL, Brothers KM, Merritt JH, Liberati NT, Ausubel FM, O’Toole GA. 2007. BifA, a cyclic-Di-GMP phosphodiesterase, inversely regulates biofilm formation and swarming motility by Pseudomonas aeruginosa PA14. J Bacteriol 189:8165–78.

53. Bassis CM, Visick KL. 2010. The cyclic-di-GMP phosphodiesterase BinA negatively regulates cellulose-containing biofilms in Vibrio fischeri. J Bacteriol 192:1269–78.

54. Rodrigues C, Passet V, Rakotondrasoa A, Brisse S. 2018. Identification of Klebsiella pneumoniae, Klebsiella quasipneumoniae, Klebsiella variicola and Related Phylogroups by MALDI-TOF Mass Spectrometry. Front Microbiol 9:3000.

55. Clegg S, Wilson J, Johnson J. 2011. More than one way to control hair growth: regulatory mechanisms in enterobacteria that affect fimbriae assembled by the chaperone/usher pathway. J Bacteriol 193:2081–8.

56. Tamayo R, Pratt JT, Camilli A. 2007. Roles of cyclic diguanylate in the regulation of bacterial pathogenesis. Annu Rev Microbiol 61:131–48.

57. Blomfield IC. 2001. The regulation of pap and type 1 fimbriation in Escherichia coli. Adv Microb Physiol 45:1–49.

58. Gally DL, Leathart J, Blomfield IC. 1996. Interaction of FimB and FimE with the fim switch that controls the phase variation of type 1 fimbriae in Escherichia coli K-12. Mol Microbiol 21:725–38.

59. Romling U, Simm R. 2009. Prevailing concepts of c-di-GMP signaling. Contrib Microbiol 16:161–181.

60. Romling U, Amikam D. 2006. Cyclic di-GMP as a second messenger. Curr Opin Microbiol 9:218–28.

61. Schmidt AJ, Ryjenkov DA, Gomelsky M. 2005. The ubiquitous protein domain EAL is a cyclic diguanylate-specific phosphodiesterase: enzymatically active and inactive EAL domains. J Bacteriol 187:4774–81.

62. Tamayo R, Tischler AD, Camilli A. 2005. The EAL domain protein VieA is a cyclic diguanylate phosphodiesterase. J Biol Chem 280:33324–30.

63. Ares MA, Fernandez-Vazquez JL, Pacheco S, Martinez-Santos VI, Jarillo-Quijada MD, Torres J, Alcantar-Curiel MD, Gonzalez YMJA, De la Cruz MA. 2017. Additional regulatory activities of MrkH for the transcriptional expression of the Klebsiella pneumoniae mrk genes: Antagonist of H-NS and repressor. PLoS One 12:e0173285.

64. Olsen PB, Klemm P. 1994. Localization of promoters in the fim gene cluster and the effect of H-NS on the transcription of fimB and fimE. FEMS Microbiol Lett 116:95–100.

65. Blomfield IC, Calie PJ, Eberhardt KJ, McClain MS, Eisenstein BI. 1993. Lrp stimulates phase variation of type 1 fimbriation in Escherichia coli K-12. J Bacteriol 175:27–36.

66. Corcoran CP, Dorman CJ. 2009. DNA relaxation-dependent phase biasing of the fim genetic switch in Escherichia coli depends on the interplay of H-NS, IHF and LRP. Mol Microbiol 74:1071–82.

67. Lang B, Blot N, Bouffartigues E, Buckle M, Geertz M, Gualerzi CO, Mavathur R, Muskhelishvili G, Pon CL, Rimsky S, Stella S, Babu MM, Travers A. 2007. High-affinity DNA binding sites for H-NS provide a molecular basis for selective silencing within proteobacterial genomes. Nucleic Acids Res 35:6330–7.

68. Olsen PB, Schembri MA, Gally DL, Klemm P. 1998. Differential temperature modulation by H-NS of the fimB and fimE recombinase genes which control the orientation of the type 1 fimbrial phase switch. FEMS Microbiol Lett 162:17–23.

69. Donato GM, Lelivelt MJ, Kawula TH. 1997. Promoter-specific repression of fimB expression by the Escherichia coli nucleoid-associated protein H-NS. J Bacteriol 179:6618–25.

70. Donato GM, Kawula TH. 1999. Phenotypic analysis of random hns mutations differentiate DNA-binding activity from properties of fimA promoter inversion modulation and bacterial motility. J Bacteriol 181:941–8.

71. Schwan WR, Lee JL, Lenard FA, Matthews BT, Beck MT. 2002. Osmolarity and pH growth conditions regulate fim gene transcription and type 1 pilus expression in uropathogenic Escherichia coli. Infect Immun 70:1391–402.

72. Ares MA, Fernandez-Vazquez JL, Rosales-Reyes R, Jarillo-Quijada MD, von Bargen K, Torres J, Gonzalez-y-Merchand JA, Alcantar-Curiel MD, De la Cruz MA. 2016. H-NS Nucleoid Protein Controls Virulence Features of Klebsiella pneumoniae by Regulating the Expression of Type 3 Pili and the Capsule Polysaccharide. Front Cell Infect Microbiol 6:13.

73. Ayala JC, Wang H, Silva AJ, Benitez JA. 2015. Repression by H-NS of genes required for the biosynthesis of the Vibrio cholerae biofilm matrix is modulated by the second messenger cyclic diguanylic acid. Mol Microbiol 97:630–45.

74. Lin TH, Chen Y, Kuo JT, Lai YC, Wu CC, Huang CF, Lin CT. 2018. Phosphorylated OmpR Is Required for Type 3 Fimbriae Expression in Klebsiella pneumoniae Under Hypertonic Conditions. Front Microbiol 9:2405.

75. Huang TW, Lam I, Chang HY, Tsai SF, Palsson BO, Charusanti P. 2014. Capsule deletion via a lambda-Red knockout system perturbs biofilm formation and fimbriae expression in Klebsiella pneumoniae MGH 78578. BMC Res Notes 7:13.

76. Datsenko KA, Wanner BL. 2000. One-step inactivation of chromosomal genes in Escherichia coli K-12 using PCR products. Proc Natl Acad Sci U S A 97:6640–5.

77. Guzman LM, Belin D, Carson MJ, Beckwith J. 1995. Tight regulation, modulation, and high-level expression by vectors containing the arabinose PBAD promoter. J Bacteriol 177:4121–30.

78. Gomes AEI, Stuchi LP, Siqueira NMG, Henrique JB, Vicentini R, Ribeiro ML, Darrieux M, Ferraz LFC. 2018. Selection and validation of reference genes for gene expression studies in Klebsiella pneumoniae using Reverse Transcription Quantitative real-time PCR. Sci Rep 8:9001.

79. Livak KJ, Schmittgen TD. 2001. Analysis of relative gene expression data using real-time quantitative PCR and the 2(-Delta Delta C(T)) Method. Methods 25:402–8.

80. Tsai KW, Lai HT, Tsai TC, Wu YC, Yang YT, Chen KY, Chen CM, Li YS, Chen CN. 2009. Difference in the regulation of IL-8 expression induced by uropathogenic E. coli between two kinds of urinary tract epithelial cells. J Biomed Sci 16:91.

81. Merritt JH, Kadouri DE, O’Toole GA. 2005. Growing and analyzing static biofilms. Curr Protoc Microbiol Chapter 1:Unit 1B 1.

82. Schwan WR, Beck MT, Hung CS, Hultgren SJ. 2018. Differential Regulation of Escherichia coli fim Genes following Binding to Mannose Receptors. J Pathog 2018:2897581.

83. Schembri MA, Blom J, Krogfelt KA, Klemm P. 2005. Capsule and fimbria interaction in Klebsiella pneumoniae. Infect Immun 73:4626–33.

84. Mulvey MA, Schilling JD, Hultgren SJ. 2001. Establishment of a persistent Escherichia coli reservoir during the acute phase of a bladder infection. Infect Immun 69:4572–9.

