## Supplemental material for "Conserved FimK truncation coincides with increased expression of type 3 fimbriae and cultured bladder epithelial cell association in *Klebsiella quasipneumoniae*"

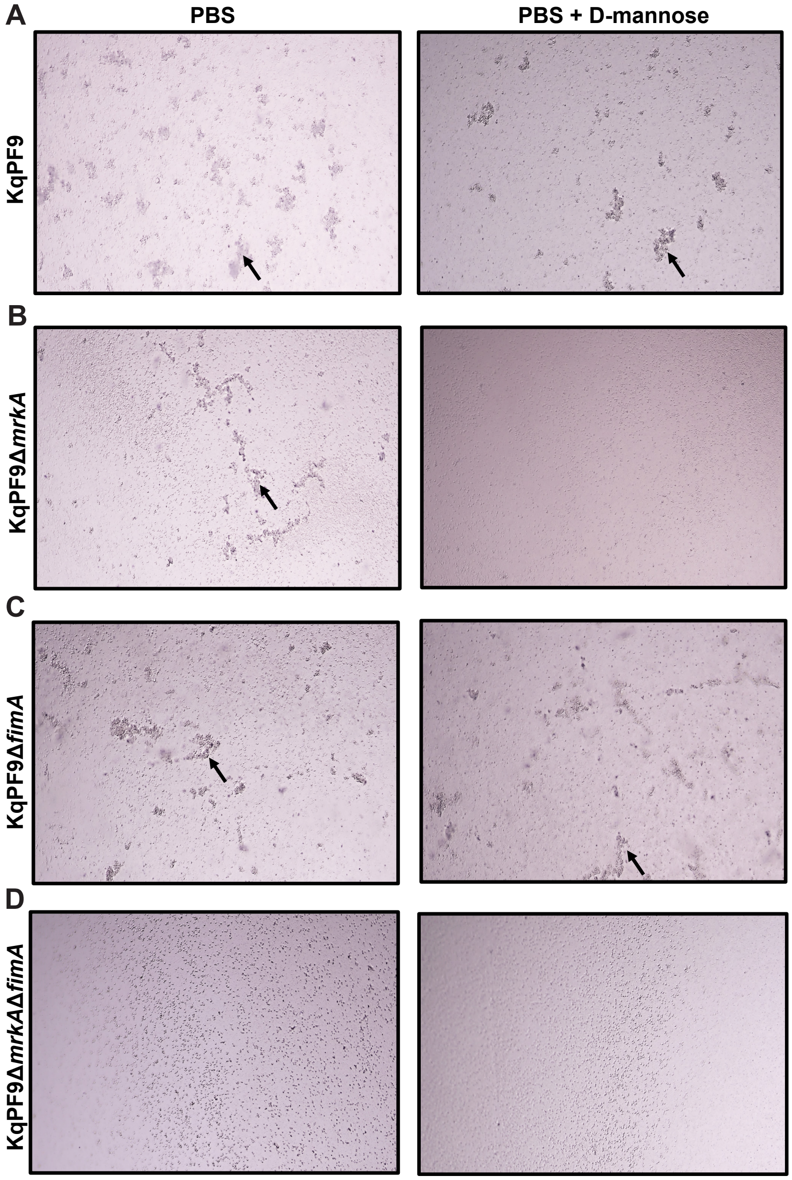


**Figure S1. KqPF9 agglutination of yeast is mannose insensitive.** Yeast agglutination (as indicated by arrows) in the presence and absence of 2.5% D-Mannose by wild-type KqPF9 and type 1 and type 3 fimbriae mutants. Yeast agglutination by **A)** KqPF9, **B)** type 3 fimbrial mutant KqPF9Δ*mrkA*, **C)** type 1 fimbrial mutant KqPF9Δ*fimA,* and **D)** KqPF9Δ*mrkA*Δ*fimA* in the absence (left) or presence (right) of 2.5% D-Mannose. KqPF9Δ*mrkA* cannot agglutinate yeast in the presence of mannose and KqPF9Δ*mrkA*Δ*fimA* cannot agglutinate yeast in the presence or absence of 2.5% D-mannose. Images were taken with a 10X objective and are representative of experiments conducted in biological triplicate.

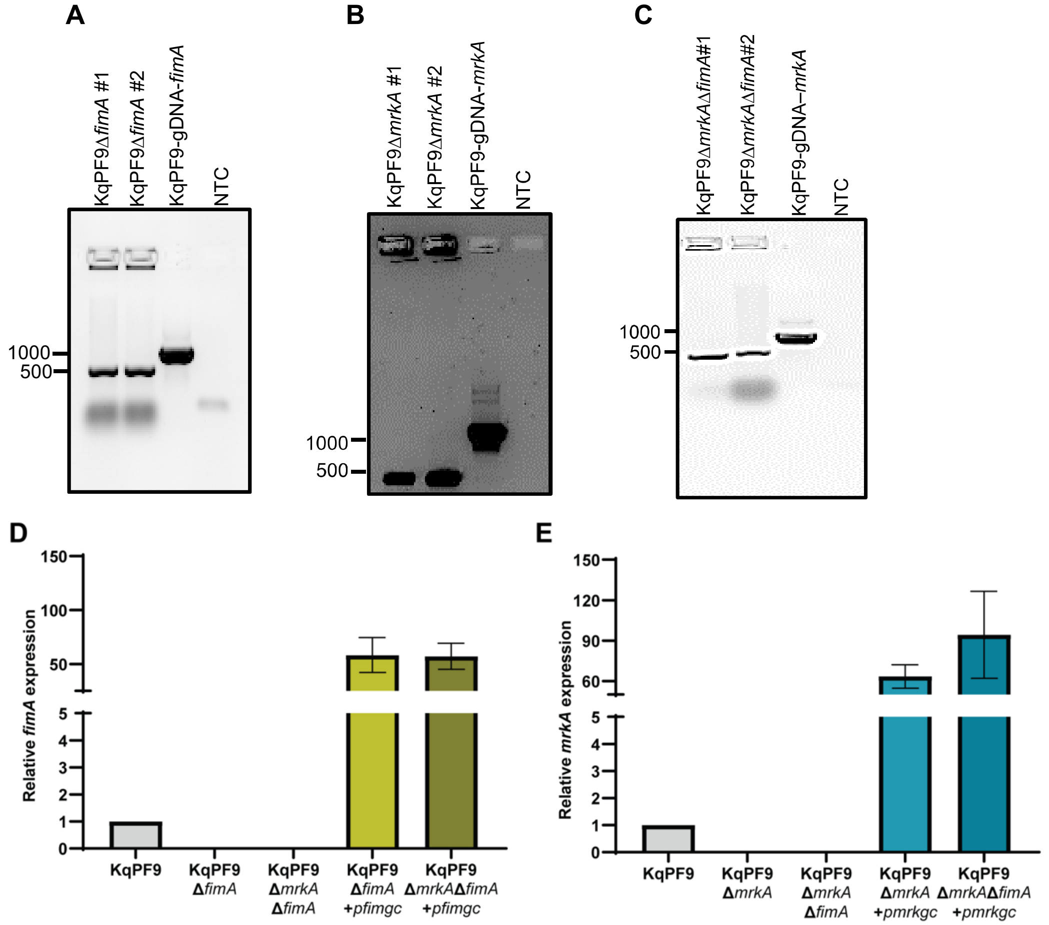


**Figure S2. KqPF9 type 1 and type 3 fimbriae knockout and complement confirmation.** Images of the PCR products indicating successful gene deletion in **A)** type 1 (KqPF9Δ*fimA*), **B)** type 3 (KqPF9Δ*mrkA*) or **C)** type 1 and type 3 fimbriae (KqPF9Δ*mrkA*Δ*fimA*) double mutants of KqPF9 resolved on an 1% agarose gel. Wildtype KqPF9 was used as positive control to verify PCR product size and water as a no template control (NTC). **D)** Quantitative reverse transcriptase PCR (qRT-PCR) analysis of type 1 fimbriae (*fimA*) expression in KqPF9 and respective isogenic mutant and complement strains. *fimgc* indicates the *fim* gene cluster (*fimAICDFGHK*). **E)** qRT-PCR analysis of type 3 fimbriae (*mrkA*) expression in KqPF9 wild-type and respective isogenic mutants and complement strains. *mrkgc* indicates the *mrk* gene cluster (*mrkABCDF*). The expression of each gene was normalized to expression of *rho* and fold change is expressed relative to wild type. Error bars indicate standard error of the mean.

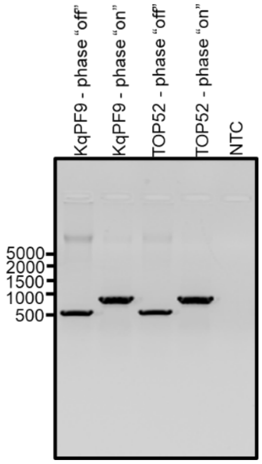


**Figure S3. *fimS* phase assay in KqPF9 and TOP52 static culture.** PCR amplification products of genomic DNA extracted from overnight static cultures of wildtype KqPF9 and TOP52 using either phase “on” or phase “off” primer sets” resolved on a 1% agarose gel. Water was used as a no template control (NTC). Banding patterns indicate *fimS* exists in both “on” and “off” orientations in KqPF9 and TOP52 during static culture.

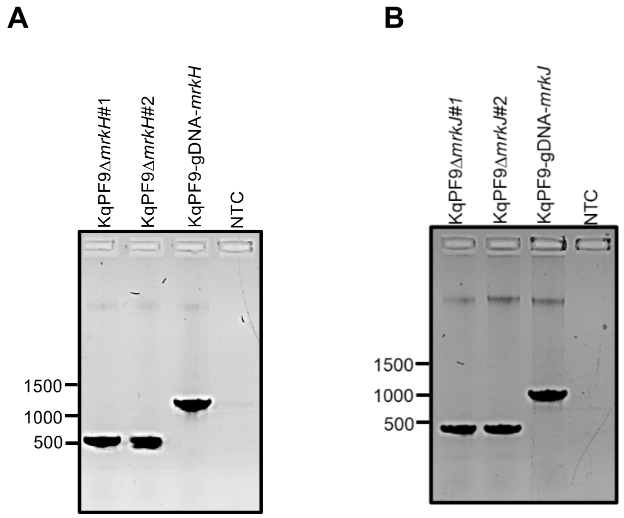


**Figure S4. Generation of isogenic *mrkH* and *mrkJ* mutants of KqPF9.** Verification PCR of isogenic **A)** *mrkH* and **B)** *mrkJ* mutants of KqPF9 resolved in an 1% agarose gel. The genomic DNA of KqPF9 was used as positive control for wildtype PCR product size and water was used as no template control (NTC).

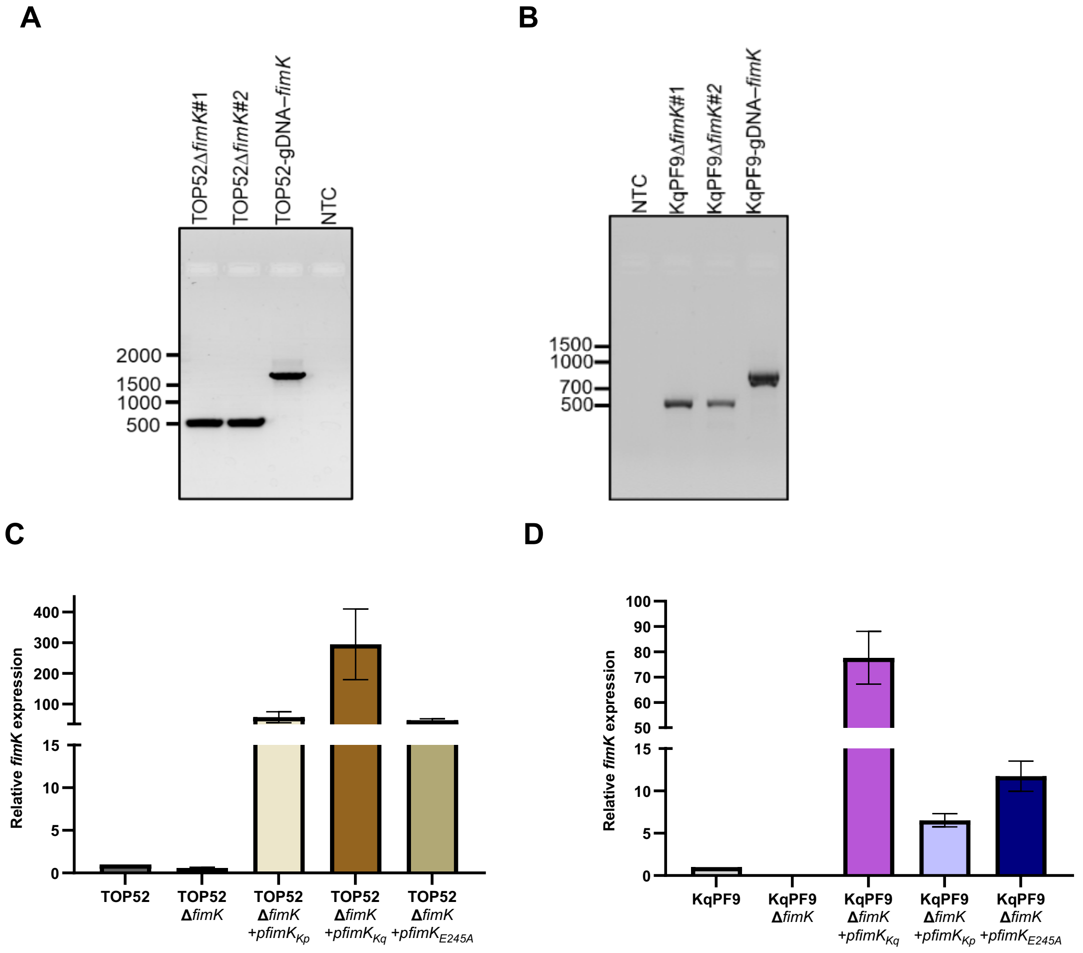


**Figure S5. Verification of isogenic *fimK* mutant and complement strains of TOP52 and KqPF9.** PCR confirmation of *fimK* knockouts in **A)** TOP52 and **B)** KqPF9 is shown as resolved through 1% agarose gel. Genomic DNA of respective wildtype strains of TOP52 and KqPF9 were used as controls to verify the PCR product size and water as a no template control (NTC). qRT-PCR analysis of *fimK* expression in **C)** TOP52 and **D)** KqPF9 and isogenic *fimK* mutants and complement strains carried out in biological and technical triplicate. *rho* was used for normalization and relative expression is in respect to wildtype. The bar plot depicts the mean and error bars indicate SEM.

**Table S1.** Primer sequences used to generate and verify isogenic mutant strains of KqPF9 and TOP52.

| Primer name | Primer use | Primer Sequence (5’- 3’) |
|---|---|---|
| PF9-*mrkA*-ko-f | Knockout cassette for *mrkA* deletion in PF9 | TATGCGAATTCACAGTGTGCTCATTGATTCGTAATTCACTCTGACAAGGAAATGGCAATGTGTGTAGGCTGGAGCTGCTTC |
| PF9-*mrkA*-ko-r | Knockout cassette for *mrkA* deletion in PF9 | CGCTTTATTATTGTTATTAACTGCCCCATCGCGGGGCAGTTTTATTTTCTGACGGAATTAGCTGACATGGGAATTAGCCATGG |
| PF9-*fimA*-ko-f | Knockout cassette for *fimA* deletion in PF9 | AGGCACAACGGCTGCCAATCCGGTTCGTTATTTCGACATCGTTCAAAGGAAAACAGTATGTGTGTAGGCTGGAGCTGCTTC |
| PF9-*fimA*-ko-r | Knockout cassette for *fimA* deletion in PF9 | TGCAAAATTAAGCGCGGGCCCTCCTGGCCCGGATGGCTTCCTTGCCTGATGTTTGCCTTAGCTGACATGGGAATTAGCCATGG |
| PF9-*mrkH*-ko-f | Knockout cassette for *mrkH* deletion in PF9 | TGTTGCTATTGCTATAAGAAAAATCAAACGCCTCACGACAACTATTTACAAGGGATGCACTGTGTAGGCTGGAGCTGCTTC |
| PF9-*mrkH*-ko-r | Knockout cassette for *mrkH* deletion in PF9 | GATAGATTGAGTGACCAATGAGATTATCATTGGTGTACAGCAATATACTTTCCAAGGGTAGCTGACATGGGAATTAGCCATGG |
| PF9-*mrkJ*-ko-f | Knockout cassette for *mrkJ* deletion in PF9 | GGTAGCCTATAATTAACCTATCCTCTGCTTATTGTCTGACTAACCTCGTGAAGAGGGATATGTGTAGGCTGGAGCTGCTTC |
| PF9-*mrkJ*-ko-r | Knockout cassette for *mrkJ* deletion in PF9 | GCAACGTGGGAATTAGTTTATAATTAAGCGGGACGAAAAAGCCGGGCAAGCCCGGCTTTTGCTGACATGGGAATTAGCCATGG |
| PF9-*fimK*-ko-f | Knockout cassette for *fimK* deletion in PF9 | GGCCAGGTCACCGCCGGCAACGTGCAGTCGATCATCGGCATTACCTTTGTCTATCAATGATGTGTAGGCTGGAGCTGCTTC |
| PF9-*fimK*-ko-r | Knockout cassette for *fimK* deletion in PF9 | CGGCACCGGTGTAAACCGGTGCGCTTTTCTCTCGCCAGCGAATCCACGCCTTTAGTCACTCATCGCTTCCCCGCTGACATGGGAATTAGCCATGG |
| TOP52-*fimK*-ko-f | Knockout cassette for *fimK* deletion in TOP52 | GGCCAGGTTACCGCCGGCAACGTGCAGTCGATCATCGGCATCACCTTTGTCTATCAATGATGTGTAGGCTGGAGCTGCTTC |
| TOP52-*fimK*-ko-r | Knockout cassette for *fimK* deletion in TOP52 | GACGATATTCGCGCATGACGTACCGGCACCGGTGCTAACCGGTGCGCTTTTCTCGCACCCGCTGACATGGGAATTAGCCATGG |
| *mrkA*-v1-for | Verify *mrkA* deletion | GCATTCTTTGACGCCGATAG |
| *mrkA*-v1-rev | Verify *mrkA* deletion | CCTGGATAAATAAAGCGGGTA |
| *fimA*-v1-for | Verify *fimA* deletion | CCACATTAAACAGATTTTAATACCG |
| *fimA*-v1-rev | Verify *fimA* deletion | CGTCGTGAAATCGGTTACTG |
| K1-rev | Kanamycin cassette | CAGTCATAGCCGAATAGCCT |

| | | |
|---|---|---|
| K2-for | Kanamycin cassette | GGTGCCCTGAATGAACTGC |
| Kt-rev | Kanamycin cassette | GGCCACAGTCGATGAATC |
| *mrkH*-v1-for | Verify *mrkH* deletion | TCCCTCCTCAATATTTGCCTG |
| *mrkH*-v1-rev | Verify *mrkH* deletion | GATTCTGATGGCAGAAATATCCT |
| *mrkJ*-v1-for | Verify *mrkJ* deletion | TTGCCGCCCTGCTCGG |
| *mrkJ*-v1-rev | Verify *mrkJ* deletion | GTTTTTACTGACGGCGGTCG |
| PF9-*fimK*-v1-for | Verify *fimK* deletion | AACGCGATCTTCACTAAC |
| PF9-*fimK*-v1-for | Verify *fimK* deletion | TGGTGGAAAAAATGCGCC |
| TOP52-*fimK*-v1-f | Verify *fimK* deletion | AACAGCACGGTCTCGCT |
| TOP52-*fimK*-v1-r | Verify *fimK* deletion | GATGGAAATACTGGAAGGG |

**Table S2.** Primer sequences used for generation of gene complements, qRT-PCR and *fimS* phase assay.

| Primer name | Primer use | Primer sequence (5'- 3') |
|---|---|---|
| *mrkABCDF*-XhoI-f | Generation of p*mrkABCDEF* | GATCCTCGAGCATGAAAAAGGTTCTTCTCTCTGCA |
| *mrkABCDF*-EcoRI-r | Generation of p*mrkABCDEF* | GATCGAATTCTTAATTATAAACTAATTCCCACGTTGC |
| *fimAICDFGHK*-XhoI-f | Generation of p*fimAICDFGHK* | GATCCTCGAGCATGAAAATCAAAACACTGGCAATG |
| *fimAICDFGHK*-EcoRI-r | Generation of p*fimAICDFGHK* | GATCGAATTCTCATGCCCGGACAAACGC |
| PF9-*fimK*-NcoI-f | Generation of p*fimK*$_{KqPF9}$ | GATCCCATGGCCGAGTACATCCTTTCT |
| PF9-*fimK*-HindIII-f | Generation of p*fimK* $_{KqPF9}$ | GATCAAGCTTTCATGCCCGGACAAACGC |
| TOP52-*fimK*-NcoI-f | Generation of *p*fimK$_{TOP52}$ | GATCCCATG GCCGATTATATCCTCTCGC |
| TOP52-*fimK*-HindIII-r | Generation of *p*fimK$_{TOP52}$ | GATCAAGCTTTCAACGTTTCGCCGGATCGC |
| TOP52-*fimK*-AIL-f | Generation of *p*fimK$_{TOP52-AIL}$ | TACAGGGGGTGGCGATCCTGATCCG |
| TOP52-*fimK*-AIL-r | Generation of *p*fimK$_{TOP52-AIL}$ | CGGATCAGGATCGCCACCCCCTGTA |
| *mrkH*-NcoI-f | Generation of p*mrkH* | GATCCCATGGCAGAGGGAACGATAAAGA |
| *mrkH*-HindIII-r | Generation of p*mrkH* | GATCAAGCTTGATTCTCTTTTTTCGCTTGGCTT |
| *mrkJ*-NcoI-f | Generation of p*mrkH* | GATCCCATGGACACTAAAATATTCGAAGACAA |
| *mrkJ*-HindIII-r | Generation of p*mrkH* | GATCAAGCTTTATGCCAATATCGTCGGCAAC |
| PF9-*mrkA*-f | qRT-PCR | GTTACCGATGTATCCTGTAC |
| PF9-*mrkA*-r | qRT-PCR | GGCAGTTAGAGACGTCAA |
| PF9-*fimA*-f | qRT-PCR | ATGATTGTTGTGTCAGCCCT |
| PF9-*fimA*-r | qRT-PCR | CCCAACTGGACGGTTTGA |
| PF9-*fimK*-f | qRT-PCR | GGTTGAGCCAGCTGATG |
| PF9-*fimK*-r | qRT-PCR | GTGGTGAGCATCCACAG |
| TOP52-*fimK*-f | qRT-PCR | GGTTGAGCCAGCTGATG |
| TOP52-*fimK*-r | qRT-PCR | GTGGTGAGCATCCACAG |
| PF9-*mrkH*-f | qRT-PCR | ATAAAATTCGCTTTCTCCTGCAT |
| PF9-*mrkH*-r | qRT-PCR | TAAACGAAAGCGGGGATCG |
| TOP52-*mrkH*-f | qRT-PCR | ATAAAATTCGCTTTCTCCTGCAT |
| TOP52-*mrkH*-r | qRT-PCR | TAAACGAAAGCGGGGATCG |
| PF9-*rho*-f | qRT-PCR | AACTACGACAAGCCGGAAAA |
| PF9-*rho*-r | qRT-PCR | ACCGTTACCACGCTCCATA |
| TOP52-*rho*-f | qRT-PCR | AACTACGACAAGCCGGAAAA |
| TOP52-*rho*-r | qRT-PCR | ACCGTTACCACGCTCCATA |
| PF9/TOP52-fimE-f | *fimS* phase "on" | GCAGGCGTATCGTATTATTCG |
| PF9/TOP52-fimS-f | *fimS* phase "off" | TGTTTTGACATATTTTGCAACTCAC |
| PF9/TOP52-fimA-r | *fimS* phase | CGTAGTAACGTGCCTGGAAC |
